# An evolutionarily conserved cis-regulatory element of *Nkx3.2* contributes to early jaw joint morphology in zebrafish

**DOI:** 10.1101/2021.11.26.470082

**Authors:** Jake Leyhr, Laura Waldmann, Beata Filipek-Górniok, Hanqing Zhang, Amin Allalou, Tatjana Haitina

## Abstract

The acquisition of movable jaws was a major event during vertebrate evolution. The role of NK3 homeobox 2 (Nkx3.2) transcription factor in patterning the primary jaw joint of gnathostomes (jawed vertebrates) is well known, however knowledge about its regulatory mechanism is lacking. In this study, we report a proximal enhancer element of *Nkx3.2* that is deeply conserved in gnathostomes but undetectable in the jawless hagfish. This enhancer is active in the developing jaw joint region of the zebrafish *Danio rerio*, and was thus designated as *jaw joint regulatory sequence 1* (JRS1). We further show that JRS1 enhancer sequences from a range of gnathostome species, including a chondrichthyan and mammals, have the same activity in the jaw joint as the native zebrafish enhancer, indicating a high degree of functional conservation despite the divergence of cartilaginous and bony fish lineages or the transition of the primary jaw joint into the middle ear of mammals. Finally, we show that deletion of JRS1 from the zebrafish genome using CRISPR/Cas9 leads to a transient jaw joint deformation and partial fusion. Emergence of this *Nkx3.2* enhancer in early gnathostomes may have contributed to the origin and shaping of the articulating surfaces of vertebrate jaws.

## Introduction

The establishment of jaw joints was one of the major events that enabled the evolutionary transition from jawless to jawed vertebrates. The earliest articulated jaws are found in fossil placoderms from the Silurian period 423 MYA (million years ago) (Zhu et al., 2013). The primary jaw joint of non-mammalian gnathostomes, including actinopterygians, amphibians, reptiles and birds is located within the first pharyngeal arch and articulates Meckel’s cartilage and the palatoquadrate. These cartilages are derived from cranial neural crest cells that migrate into the first pharyngeal arch and later ossify into anguloarticular and quadrate bones, respectively (Schilling and Kimmel, 1994; Tucker et al., 2004).

The transition from jawless to jawed vertebrates and the underlying gene regulatory network changes are not yet fully understood. However, the transcription factor Nkx3.2 (Bapx1), which acts as a chondrocyte maturation inhibitor (Provot et al., 2006), is thought to have played a major role in the evolution of the primary jaw joint (Cerny et al., 2010). *Nkx3.2* displays focal expression in the first pharyngeal arch, between Meckel’s cartilage/anguloarticular and the palatoquadrate/quadrate in non-mammalian vertebrates, as shown in zebrafish (Miller et al., 2003), *Xenopus* (Square et al., 2015), and python and chicken (Anthwal et al., 2013). In the lamprey, a jawless vertebrate, *Nkx3.2* is expressed in the ectomesenchyme surrounding the pharyngeal arches (Miyashita, 2018).

The importance of *Nkx3.2* for the development of the primary jaw joint has been shown in knock-down and knock-out experiments carried out in zebrafish and *Xenopus*. Reduction or loss of *nkx3.2* expression led to absence of the joint and fusion of Meckel’s cartilage and the palatoquadrate (Lukas and Olsson, 2018a; Miller et al., 2003; Miyashita et al., 2020; Waldmann et al., 2021). Overexpression of *nkx3.2* in *Xenopus* resulted in extra ectopic joints forming in the jaw cartilage (Lukas and Olsson, 2018b).

During the course of mammal evolution, first pharyngeal arch elements and parts of the second arch underwent a morphological transition to form the middle ear ossicles in mammals (Anthwal et al., 2013; Luo, 2007). This transition was accompanied by the development of a secondary jaw joint articulating the squamosal and dentary, which serve as the novel mandibular components. The middle ear consists of three major bones: the malleus, incus, and stapes, and the middle ear-associated bones: the gonium and tympanic ring which attach the ossicles to the skull. The incus and malleus are articulated by the incudomalleolar joint, homologous to the primary jaw joint of non-mammalian gnathostomes. *Nkx3.2* expression has been shown to be present in this joint, the tympanic ring, and the gonium, and mice homozygous for *Nkx3.2* knockouts display loss of the gonium and hypoplasia of the tympanic ring but minimal disruption to the incudomalleolar joint (Tucker et al., 2004). To what degree the gene regulatory network for these homologous joints is conserved is not fully understood.

In this study, we identified a novel gnathostome-specific cis-regulatory element, *jaw joint regulatory sequence 1* (JRS1), proximal to the *Nkx3.2* gene. We show that JRS1 has a highly conserved sequence and demonstrate that JRS1 sequences from multiple gnathostomes drive fluorescent reporter expression in the primary jaw joint of zebrafish, implying functional conservation between diverse clades. To test if JRS1 is essential for the jaw joint development, we generated a CRISPR/Cas9-induced enhancer knockout zebrafish line and show that homozygous mutants display transient dysmorphology and partial fusion of the jaw joint-articulating cartilages.

## Results

### *Nkx3.2* is located in a conserved syntenic region in vertebrates

In order to search for proximal conserved non-coding elements (CNEs), we first performed an analysis of the gene synteny around *Nkx3.2*, reasoning that if we find homologous genes upstream and downstream of *Nkx3.2* in different vertebrate species, the locus is unlikely to have undergone major rearrangements that would have disrupted the intergenic non-coding sequences. Our synteny analysis showed that *Nkx3.2* is located in a highly conserved syntenic region in all examined jawed vertebrate genomes between *Bod1l1* (Biorientation of chromosomes in cell division protein 1-like 1) and *Rab28* (Ras-related protein Rab-28) genes (Figure 1). The upstream gene *Bod1l1* and downstream gene *Rab28* have the same orientation (located on the same DNA strand) as *Nkx3.2*. However, we could not identify *Bod1l1* in the coelacanth genome or *Rab28* in the zebrafish genome. In the hagfish genome, *Nkx3.2* (ENSEBUG00000015739) is located on the contig FYBX02009947.1 next to *Nkx2.5* (ENSEBUG00000006582), but on the opposite DNA strand. *Rab28* (ENSEBUG00000007671) is located downstream from *Nkx3.2*, but there is an additional gene, annotated as Spondin 2 (*Spon2*, ENSEBUG00000016900) between them (Figure 1). These results indicated that the intergenic sequences flanking *Nkx3.2* were likely to be homologous, and therefore were appropriate to align in search of conserved non-coding elements (CNEs).

**Figure 1.**
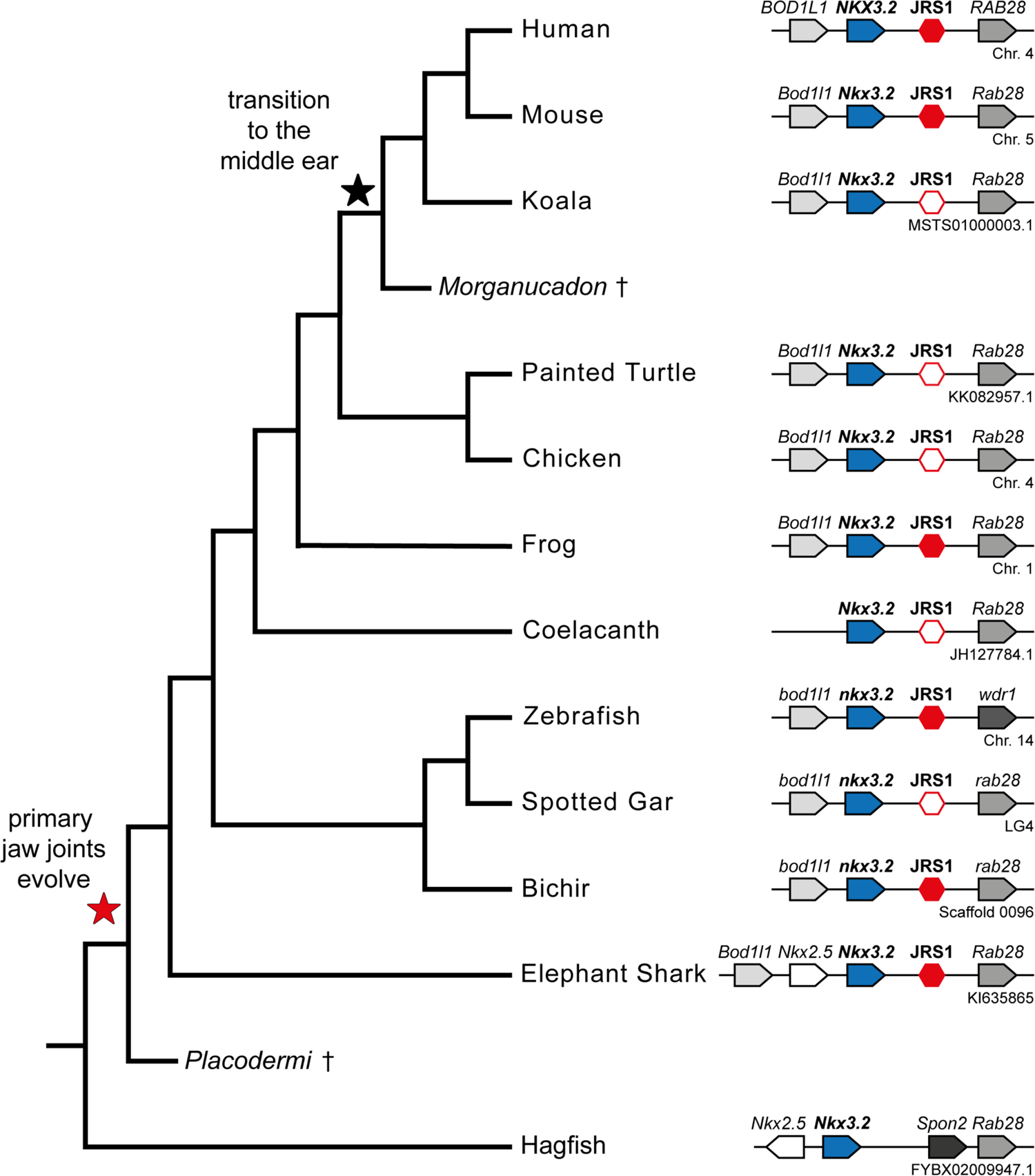
Gene synteny around *Nkx3.2* in vertebrate genomes. Phylogenetic tree based on commonly accepted topology. Pointed boxes represent gene orientation with gene names stated on top. Red hexagons mark the position of conserved non-coding element (JRS1) downstream of *Nkx3.2*, where filled hexagons mark CNEs selected for *in vivo* functional characterization in this study. The corresponding chromosome/contig number is marked below the gene order schematic of each species.

### A conserved non-coding element is identified proximal to *Nkx3.2*

mVISTA analysis identified a conserved peak in the non-coding region downstream of *Nkx3.2*, between *Nkx3.2* and *Rab28*, for all examined gnathostome species: human, mouse, koala, painted turtle, chicken, frog, coelacanth, zebrafish, spotted gar, bichir, and elephant shark (Figure 1A). We could not identify the same conserved peak downstream of *Nkx3.2* in hagfish, either upstream or downstream of *Spon2*. The peak sequences from several key gnathostome species (human, mouse, frog, zebrafish, bichir, and elephant shark) were extracted and a search for conserved motifs within these peak sequences was performed with MEME, identifying a core ∼245bp sequence (Supplementary Figure 2) containing four conserved motifs, two of which (1 and 4) were absent in the elephant shark, and one of which (4) was absent in zebrafish (Figure 2B).

**Figure 2.**
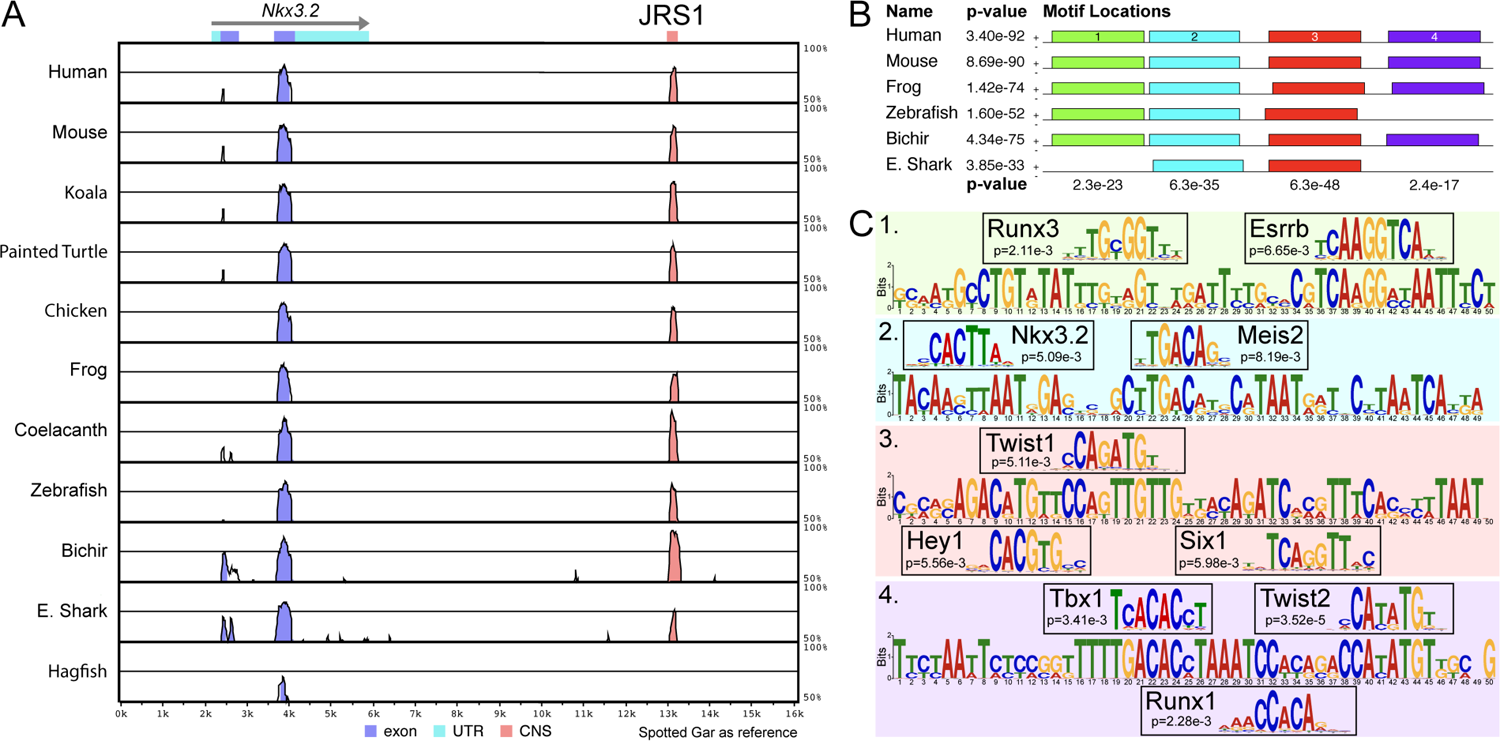
A conserved non-coding element JRS1 identified using mVISTA and MEME. (**A**) mVISTA alignment of vertebrate *Nkx3.2* loci, using the spotted gar locus as reference. Peaks indicate conserved sequences >50% identity, coloured peaks indicate >70% identity. Dark blue peaks indicate conserved exon sequences, pink indicates conserved non-coding sequences. (**B**) Shared sequence motifs (1-4) in the core ∼245bp sequence of JRS1 in different species identified with MEME analysis. P-values indicated per species (horizontal) and per motif (vertical). (**C**) Relevant transcription factor binding sites predicted by Tomtom at motifs 1-4 with associated p-values for each match.

### JRS1 revealed significant matches to transcription factors for pharyngeal arch patterning

Comparison of each 49-50bp motif within the JRS1 core sequence with known transcription factor binding motifs revealed significant matches to transcription factors known to be involved in pharyngeal arch patterning, skeletogenesis and joint formation (Figure 2C), consistent with a predicted regulatory role for JRS1 in the jaw joint expression of *Nkx3.2*.

Runx3 regulates target genes involved in chondrocyte development (Wigner et al., 2013), while Runx1 has been implicated in suppressing chondrocyte maturation by upregulating *Nkx3.2* expression (Yano et al., 2019). Hey1 is a transcriptional repressor expressed in the dorsal domain of the developing pharyngeal arches downstream of Jagged-Notch signalling, where it may serve to reinforce dorsal identity (Zuniga et al., 2010). The presence of a binding motif in JRS1 suggests that Hey1 may function in repressing dorsal expression of *Nkx3.2*, restricting expression to the intermediate domain where the jaw joint is patterned. Hey1 has also been found to be upregulated in osteoarthritis (OA) joint cartilage (Chang et al., 2021).

Tbx1 has shown to interact with Hand2 and Edn1, key regulators of pharyngeal arch development, such that zebrafish Tbx1 mutants show drastic reductions in pharyngeal cartilage (Piotrowski et al., 2003). In addition to Tbx1, the same binding region matches many other Tbx factors as they all possess highly similar binding motifs, and some of these Tbx factors are also known to function in craniofacial development (Papaioannou, 2014). For example, *tbx22* expression mirrors that of *nkx3.2* in the developing zebrafish jaw joint (Jezewski et al., 2009; Swartz et al., 2011), and although the Tbx22 binding motif is absent from the JASPAR database, the conservation of Tbx factor binding motifs suggests that Tbx22 can likely bind at this motif in JRS1, implying a possible role for Tbx22 in jaw joint formation upstream of Nkx3.2. Twist1 and Twist2 have both been shown to function in both promoting and inhibiting chondrogenesis in different contexts (Cleary et al., 2017; Reinhold et al., 2006; Takai et al., 2019), and are expressed in the zebrafish craniofacial skeleton (Germanguz and Gitelman, 2012). *Twist1* is upregulated in osteoarthritis (OA) (Hasei et al., 2017) while *Nkx3.2* is downregulated (Caron et al., 2015; Oh et al., 2021), suggesting that repression of *Nkx3.2* expression by Twist1 may contribute to OA articular cartilage pathology. *Six1* has been found to have expression restricted to articular cartilage in porcine knee joints (Hissnauer et al., 2010), and has a role in craniofacial skeletogenesis promoting Jagged-Notch signalling and repressing *Edn1* expression in the dorsal pharyngeal arches (Tavares et al., 2017). It is also upregulated by the aforementioned Tbx1 (Guo et al., 2011). Meis2 is known to function in the development of neural crest derivative tissues including cranial cartilage and bone (Machon et al., 2015), and fusions between Meckel’s cartilage and the palatoquadrate have been observed in *meis2*-knockdown zebrafish (Melvin et al., 2013).

A potential function of Esrrb in skeletogenesis has not been described, but closely-related members of the Estrogen-Related receptor family with similar binding motifs, Esrra and Esrrg, are known to function in promoting chondrogenesis or cartilage degradation in OA (Kim et al., 2015; Son et al., 2017; Tang et al., 2021). Finally, the presence of an Nkx3.2-binding motif in JRS1 suggests a potential for autoregulation of *Nkx3.2* expression, establishing a steady expression level at a reduced metabolic cost (Mcadams and Arkin, 1997).

### Acanthopterygian-specific loss, divergence, or translocation of JRS1

In addition to the broad sampling of gnathostome species, we performed a more detailed analysis of teleost fish species beyond just the zebrafish. mVISTA analysis of the non-coding region between *Nkx3.2* and *Rab28* (or *wdr1* in zebrafish) was performed for all the previously mentioned gnathostome species plus a different teleost species, one at a time. These additional teleost species included the arowana (*Sclerophages formosus*), electric eel (*Electrophorus electricus*), cavefish (*Astyanax mexicanus*), atlantic salmon (*Salmo salar*), atlantic cod (*Gadus morhua*), soldierfish (*Myripristis murdjan*), seahorse (*Hippocampus comes*), tilapia (*Oreochromis niloticus*), amazon molly (*Poecilia formosa*), stickleback (*Gasterosteus aculeatus*), and pufferfish (*Tetraodon nigroviridis*). We were able to identify JRS1 in all teleost species with the exception of the soldierfish, seahorse, tilapia, amazon molly, stickleback, and pufferfish - all members of the clade Acanthopterygii (Hughes et al., 2018), suggestive of an acanthopterygian-specific loss, divergence, or translocation of JRS1 (Supplementary Figure 1).

### Generation of *nkx3.2(*JRS1):mCherry transgenic lines

To test JRS1 for enhancer activity *in vivo*, we generated Tol2 reporter constructs with species-specific JRS1 sequences from human, mouse, frog, zebrafish, bichir, and elephant shark upstream of a membrane-tagged mCherry coding sequence. Embryos injected with JRS1 reporter constructs displayed mosaic reporter gene expression within the first pharyngeal arch elements Meckel’s cartilage and palatoquadrate. At least two positive founders were identified for each injected construct and used for generation of stable *nkx3.2*(JRS1):mCherry transgenic lines, which allowed further characterization of enhancer activity.

### Zebrafish JRS1 enhancer drives fluorescent reporter gene expression corresponding to endogenous *nkx3.2* expression

For analysing zebrafish JRS1 activity in the transgenic fluorescent reporter line, we performed live confocal imaging at different developmental time-points. From 40 hours post fertilization (hpf) we detected persistent mCherry expression driven by the *nkx3.2* enhancer within the jaw joint-forming region of the first pharyngeal arch (Figure 3, Video 1). At 30 hours post fertilization the *nkx3.2*(JRS1):mCherry line furthermore displayed mCherry expressing cells in the notochord, which could be detected up to 3 days post fertilization (dpf) (Video 1).

**Figure 3.**
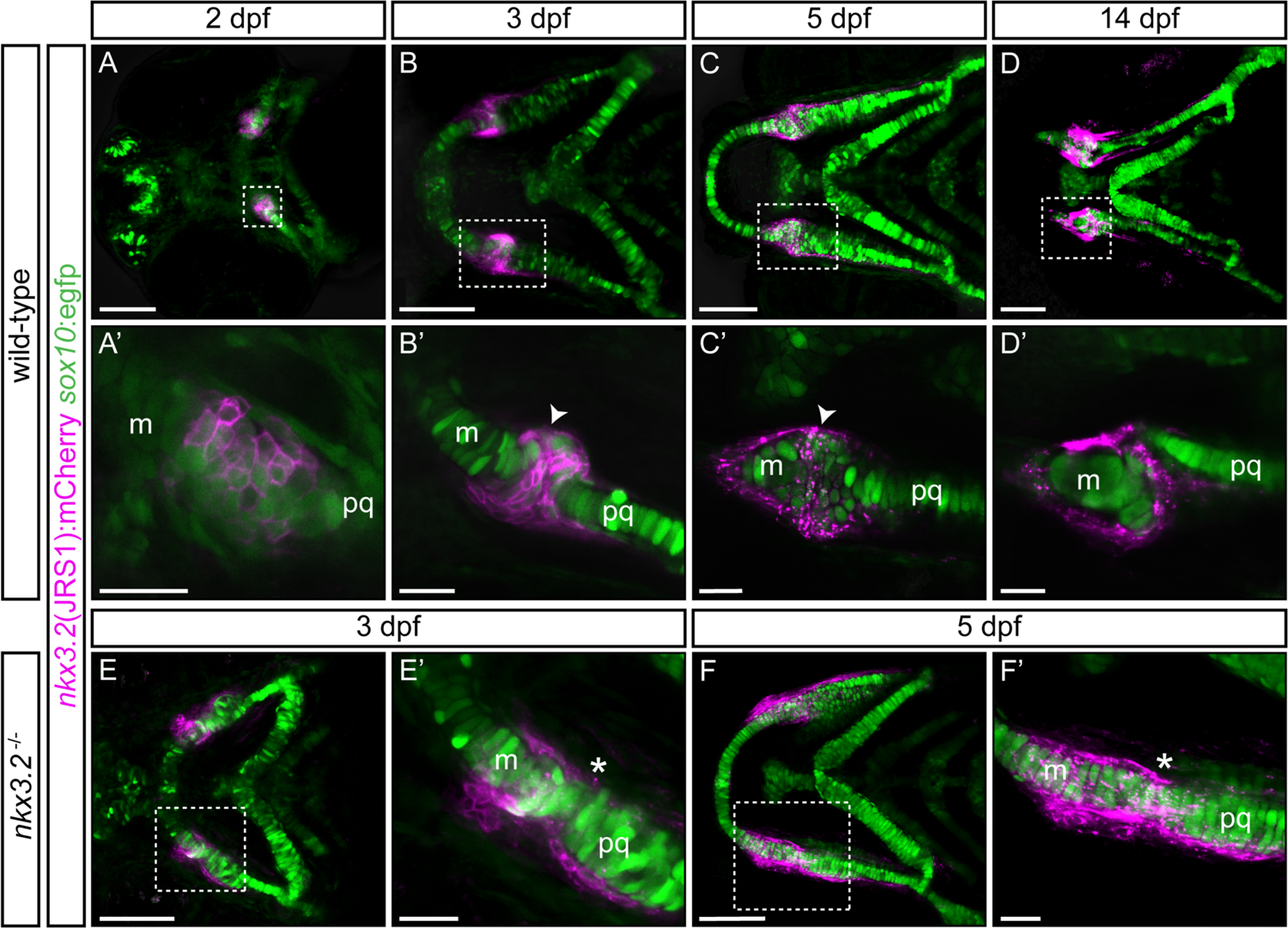
Zebrafish JRS1 enhancer drives mCherry reporter gene expression in jaw joint-forming chondroprogenitor cells and partly overlaps with *sox10:*egfp expressing cells. (**A, A’**) At 2 dpf jaw joint progenitor cells express both GFP and mCherry. (**B, B’)** By 3 dpf cells start to differentiate and *nkx3.2*(JRS1):mCherry activity is restricted to jaw joint-forming interzone, overlapping with *sox10*:egfp. Single-labelled mCherry expressing cells are surrounding the joint-forming region. (**C, C’)** At 5 dpf mCherry expressing cells are restricted to the articulation forming area between Meckel’s cartilage (m) and the palatoquadrate (pq). Double mCherry/GFP expressing cells are restricted to posterior Meckel’s cartilage and anterior palatoquadrate. (**D, D’)** At 14 dpf a clear joint cavity is visible. *nkx3.2*(JRS1):mCherry activity is restricted to the joint cavity and to both lateral and medial palatoquadrate. Dashed box in **A-D** is magnified in **A’-D’**. **A-D** represents maximum projection of confocal Z-stack, and **A’-D’** represents a single confocal image. In *nkx3.2*^-/-^ mutants *nkx3.2*(JRS1):mCherry marks the cells outside of the fused jaw joint at 3dpf (E, E’) and 5 dpf (F, F’). Dashed box in **E-F** is magnified in **E’-F’**. **E-F** and **E’-F’** represent maximum projection of confocal Z-stack. Scale bars: 100 μm **(A-F)** and 25 μm **(A’-F’)**.

**Video 1 | JRS1 enhancer drives reporter expression in the early notochord and jaw joints.** Lateral view of *nkx3.2*(JRS1):mCherry/*sox10:*egfp zebrafish embryo developing from 44-71 hpf. Inset *in situ* hybridisation image of *nkx3.2* expression in 2 dpf zebrafish embryo. White arrowhead indicates jaw joint expression domain while black arrowheads indicate expression in the notochord.

*nkx3.2*(JRS1):mCherry/*sox10*:egfp double transgenic fish were used for the characterization of joint progenitor cells from 2 dpf. The *sox10*:egfp line labels neural crest cell derived populations comprising pharyngeal arch cartilages (Carney et al., 2006). At the onset of chondrogenesis (2 dpf), confocal live imaging revealed overlapping expression of *nkx3.2*(JRS1):mCherry and *sox10*:egfp in condensed mesenchyme cells in the jaw joint establishing zone (Figure 3A, A’). Single *sox10:*egfp expressing cells were located adjacent to this area, labelling first pharyngeal arch forming elements Meckel’s cartilage and palatoquadrate (Figure 3A’). All labelled cells displayed rounded pentagon morphology at this stage.

By 3 dpf *nkx3.2*(JRS1):mCherry expressing cells were densely packed in the joint-forming region, overlapping with *sox10:*egfp expressing cells (Figure 3B, B’) and displayed more elongated morphology. In Meckel’s cartilage and the palatoquadrate, the majority of GFP-labelled chondrocytes underwent differentiation and maturation indicated by the “coin stack” arrangement (Figure 3B, B’). Double-labelled mCherry/GFP cells were furthermore present in the developing retroarticular process (RAP) (Figure 3B, B’). At both medial and lateral surfaces of the joint, single *nkx3.2*(JRS1):mCherry expressing cells were lined up from the posterior Meckel’s cartilage up to the anterior palatoquadrate.

mCherry expression became increasingly scattered in the membrane of labelled cells by 5 dpf. We observed a consistent presence of mCherry-positive cells on the lateral edge of the palatoquadrate and posterior Meckel’s cartilage, partly overlapping with GFP-positive cells (Figure 3C, C’). The morphology of double-labelled cells in and surrounding the jaw joint-forming region was distinct from chondrogenic cells forming the main cartilage elements, by displaying smaller size and more rounded morphology (Figure 3C,C’). By 14 dpf, the joint cavity was evident and contained *nkx3.2*(JRS1):mCherry expressing cells (Figure 3D,D’). mCherry-positive cells were maintained in the posterior part of Meckel’s cartilage and we observed a line of mCherry-positive cells on both the lateral and medial sides of the palatoquadrate, reminiscent of the perichondrium, extending posteriorly towards the hyosymplectic of the second pharyngeal arch (Figure 3D,D’).

In order to characterize the *nkx3.2*(JRS1):mCherry expressing cells in *nkx3.2* gene mutants we incrossed previously reported *nkx3.2^+/uu2803^* mutant alleles (Waldmann et al., 2021) with *nkx3.2*(JRS1):mCherry/*sox10*:egfp background and used these heterozygotes to generate homozygous *nkx3.2^uu2803/uu2803^* mutants (abbreviated to *nkx3.2*^-/-^). Notably, *nkx3.2*(JRS1):mCherry expressing cells were detected lining the outside of the cartilage in the fused jaw joint region of *nkx3.2*^-/-^ mutants at 3-5 dpf (Figure 3E-F, E’-F’). The presence of double labelled cells was not as apparent as in wild-type fishes, however the scattering of the membranal mCherry signal was also noticeable in *nkx3.2*^-/-^ mutants from 5 dpf (Figure 3F, F’).

### JRS1 activity is conserved in a range of gnathostome species

Generated transgenic lines containing JRS1 enhancer sequences from human, *Homo sapiens* (Figure 4A); mouse, *Mus musculus* (Figure 4B); frog, *Xenopus tropicalis* (Figure 4C); bichir, *Polypterus senegalus* (Figure 4H); and elephant shark, *Callorhinchus milii* (Figure 4I) displayed mCherry reporter activity in the developing jaw joint (Figure 4), consistent with the zebrafish JRS1 activity (Figure 4G). In contrast to zebrafish JRS1 however, early (∼30 hpf) mCherry expression in the notochord could not be detected in these lines. *fli1a:gfp* was used as background marker in order to investigate the potential JRS1 activity in the perichondrium. mCherry expression in perichondrium cells surrounding the developing jaw joint and palatoquadrate was present in all tested *nkx3.2*(JRS1):mCherry enhancer lines (Figure 4D-F’; J-L’).

**Figure 4.**
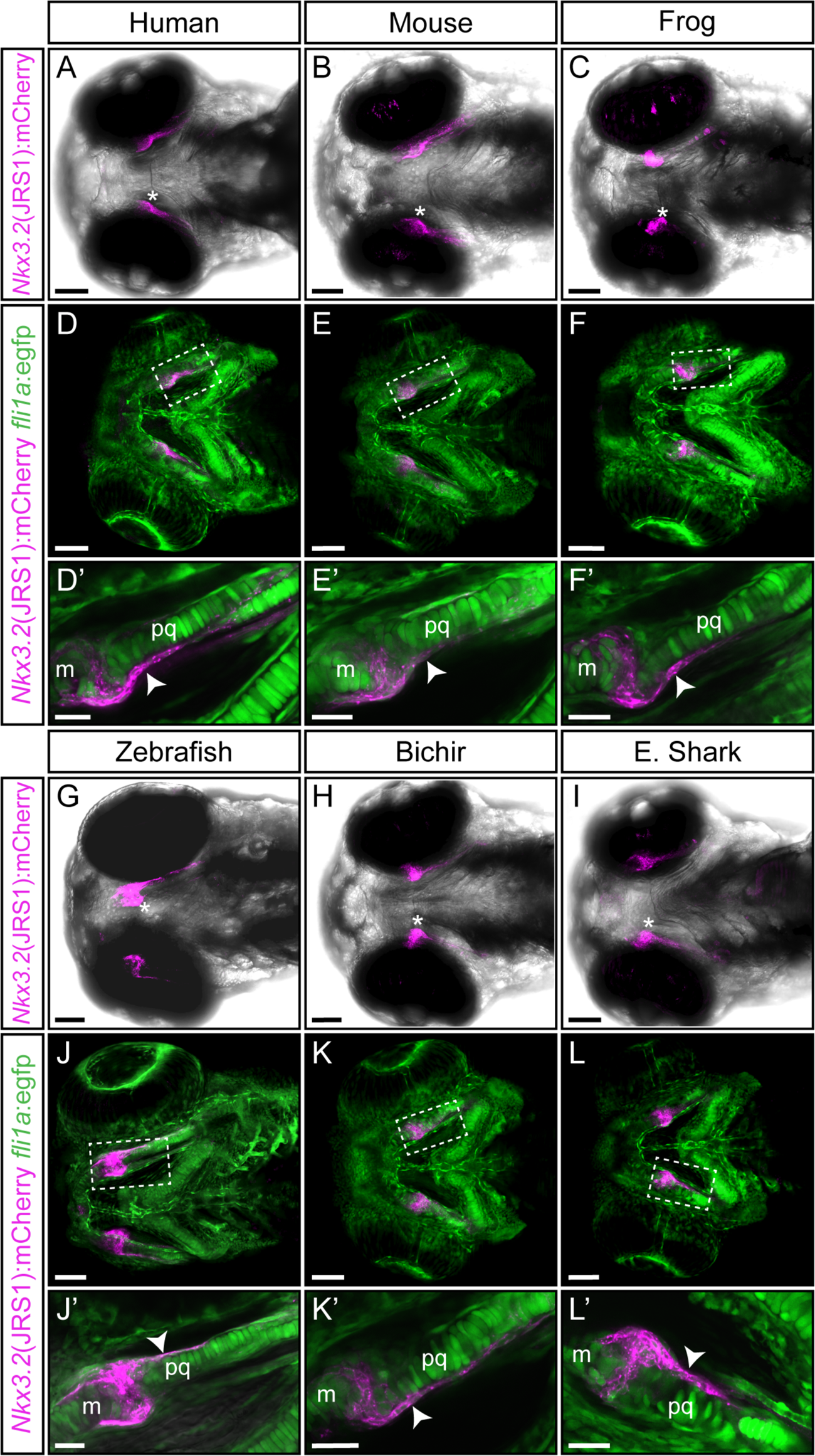
Functional conservation of the JRS1 enhancer within tested gnathostome species. (**A-C, G-I**) Maximum projection images of 3 dpf transgenic zebrafish embryos (ventral view) driving mCherry expression in jaw joint and mandibular arch elements under the control of the JRS1 sequence of (**A**) human *Homo sapiens*, (**B**) mouse *Mus musculus*, (**C**) frog *Xenopus tropicalis*, (**G**) zebrafish *Danio rerio,* (**H**) bichir *Polypterus senegalus* and (**I**) elephant shark *Callorhinchus milii*. Asterisk marks jaw joint. (**D-F**; **J-L**) Maximum projection images of 3 dpf *nkx3.2*(JRS1):mCherry transgenic zebrafish driving mCherry expression under the control of species-specific enhancer sequences with *fli1a*:egfp background reveals mCherry expression in GFP-labelled perichondrium cells. Dashed box is magnified in **D’-F’** and **J’-L’** as single confocal images. White arrowhead marks mCherry expression in the perichondrium. m: Meckel’s cartilage; pq: palatoquadrate. Scale bars: 75 μm (**A-L**), 25 μm (**D’-F’** and **J’-L’**).

### JRS1 deletion results in transient jaw joint dysmorphology and partial fusion

As JRS1 appears functionally conserved and active specifically in the jaw joint, we next investigated the function more closely by using CRISPR/Cas9 genome editing to generate a zebrafish line with the entire JRS1 enhancer sequence deleted from the genome. Two sgRNAs were targeted to sequences either side of JRS1 resulting in a 445bp deletion spanning the conserved core of the JRS1 and ∼100 bp flanking sequences (Figure 5A, Supplementary Figure 3). The resulting enhancer deletion allele is termed *nkx3.2*^ΔJRS1^. We generated homozygous mutant zebrafish (*nkx3.2*^⊗JRS1/⊗JRS1^) and used Alcian blue and alizarin red staining to characterise the craniofacial morphology in comparison to heterozygotes and wild-type siblings at 5, 9, 14, and 30 dpf.

**Figure 5.**
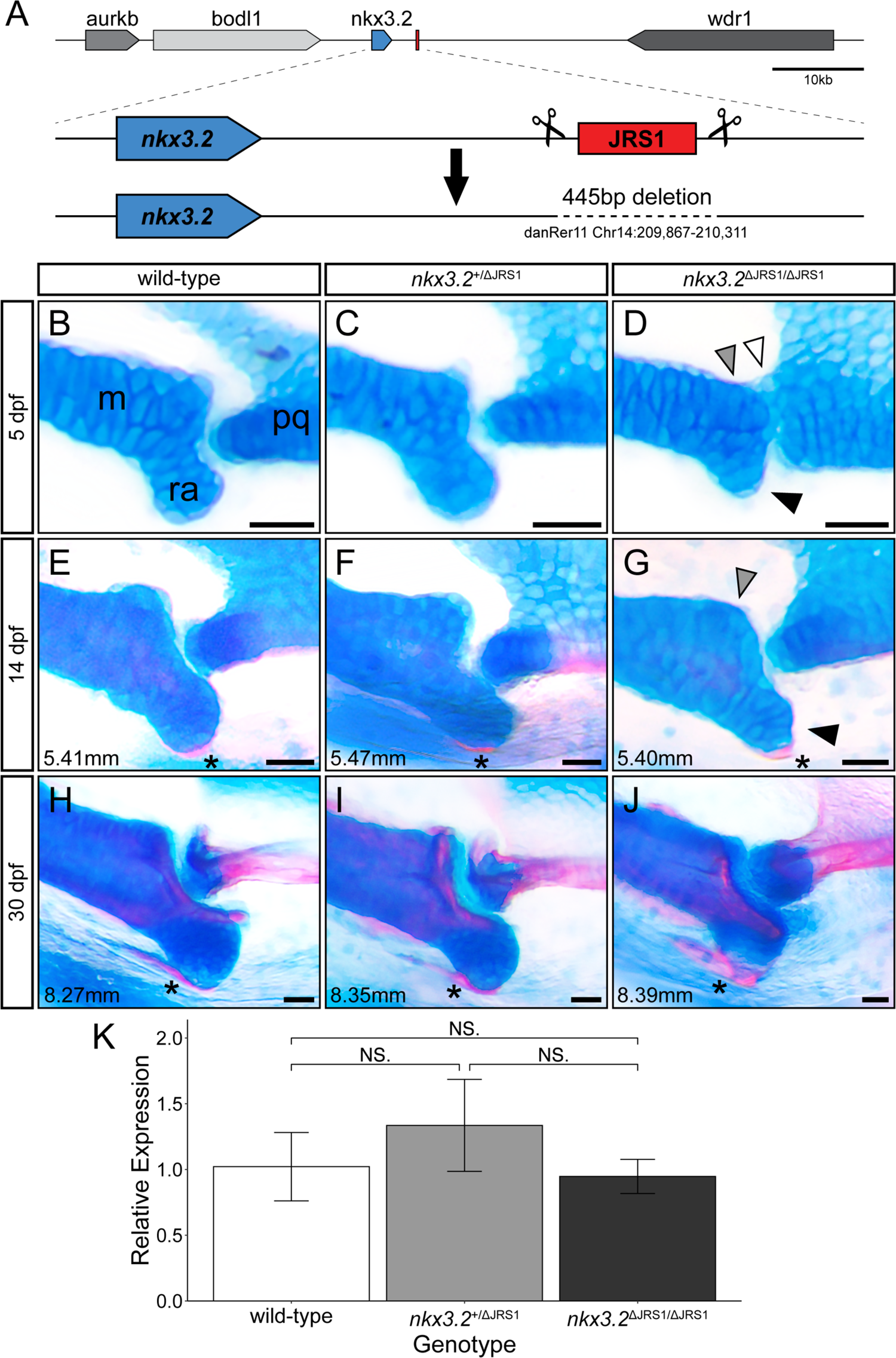
Homozygous JRS1 enhancer deletion results in early jaw joint dysmorphology. (**A**) Schematic of the JRS1 deletion allele generation. (**B-J**) Alcian blue and alizarin red-stained jaw joints of wild-type, *nkx3.2*^+/_Δ_JRS1^, and *nkx3.2*^_Δ_JRS1/_Δ_JRS1^ zebrafish at 5, 14, and 30 dpf. Standard lengths (mm) are given for 14 dpf and 30dpf juveniles. White arrowhead marks the partial fusion between Meckel’s cartilage (m) and the palatoquadrate (pq). Grey arrowhead indicates the rounded, non-bulbous anteroposterior process of Meckel’s cartilage. Black arrowhead marks the reduced retroarticular process (ra). Asterisx marks the ossifying retroarticular bone. Scale bars: 25 µm. (**K**) Relative *nkx3.2* expression levels at 6 dpf in wild-type, *nkx3.2*^+/_Δ_JRS1^, and *nkx3.2*^_Δ_JRS1/_Δ_JRS1^ zebrafish. NS. denotes P>0.05 (wild-type vs *nkx3.2*^_Δ_JRS1/_Δ_JRS1^ P=1.0, *nkx3.2*^+/_Δ_JRS1^ vs wild-type and *nkx3.2*^_Δ_JRS1/_Δ_JRS1^ both P=0.6).

At all examined ages, the jaw joint morphology of wild-type and heterozygous mutants appeared indistinguishable. At 5 dpf, homozygous *nkx3.2*^_Δ_JRS1/_Δ_JRS1^ mutants tended to display a reduced RAP on Meckel’s cartilage and the cartilage of the palatoquadrate often failed to form a pronounced convex joint process (Figure 5D). The result of this was a lack of a clearly defined concave-convex articulation in the jaw joint. Additionally, while no mutants displayed a complete fusion between the Meckel’s and palatoquadrate cartilages, there was often a small number of chondrocytes (white arrowhead, Figure 5D) connecting the two elements. Mouth gape angle did not appear to be affected, suggesting that the joint was at least partially flexible at this stage.

At 14 dpf, the RAP of homozygous mutants became more pronounced and comparable to wild-type and heterozygous siblings (Figure 5G). Meckel’s cartilage and the palatoquadrate appeared to be fully separated, and the palatoquadrate joint process was more convex, resulting in a more concave-convex shape to the joint articulation. The RAP began to ossify as normal in all genotypes. By 30 dpf, the jaw joint of homozygous mutants was indistinguishable from wild-type and heterozygote siblings, with ossification of Meckel’s cartilage, the RAP, and the palatoquadrate progressing normally (Figure 5J).

The relative expression levels of *nkx3.2* were quantified in JRS1 mutants using qPCR to determine the contribution of JRS1 to gene expression at 6 dpf. Larvae were individually genotyped using tail tissue and then pooled into 3 biological replicates of 7 larvae per genotype. Following RNA extraction and cDNA synthesis, qPCR was performed and *nkx3.2* expression of heterozygous and homozygous JRS1 mutants was compared relative to wild-type. No significant differences between the genotypes were detected at this time point (Figure 5K).

To quantify the subtle differences in posterior Meckel’s cartilage shape in JRS1 mutant larvae, the heads 9 dpf *nkx3.2*^+/+^ (n=12), *nkx3.2*^+/_Δ_JRS1^ (n=10), and *nkx3.2*^_Δ_JRS1/_Δ_JRS1^ (n=8), were imaged using OPT, and the whole heads (Supplementary Figure 4) and both jaw joints were rendered as maximum projections for a total of 60 jaw joints. The shape of Meckel’s cartilage at the joint interface was analysed using 2D geometric morphometrics (Figure 6A), confirming the previously described observations that there was no significant difference between wild-type and heterozygous mutants (P=0.20) while homozygous mutants differed significantly compared to them both (P=0.0015). Analysis of morphospace also confirmed the tendency for homozygous mutants to display a reduced RAP resulting in a less concave surface interfacing with the palatoquadrate (Figure 6A). This phenotype is not fully penetrant, as there is some overlap between the range of wild-type and mutant shapes, consistent with the variable severity of mutant joint phenotypes seen at 5 dpf. Averaged 3D renderings of the jaw joint at 9 dpf also displayed the partial fusion between Meckel’s cartilage and the palatoquadrate previously described at 5 dpf (Figure 6D, G).

**Figure 6.**
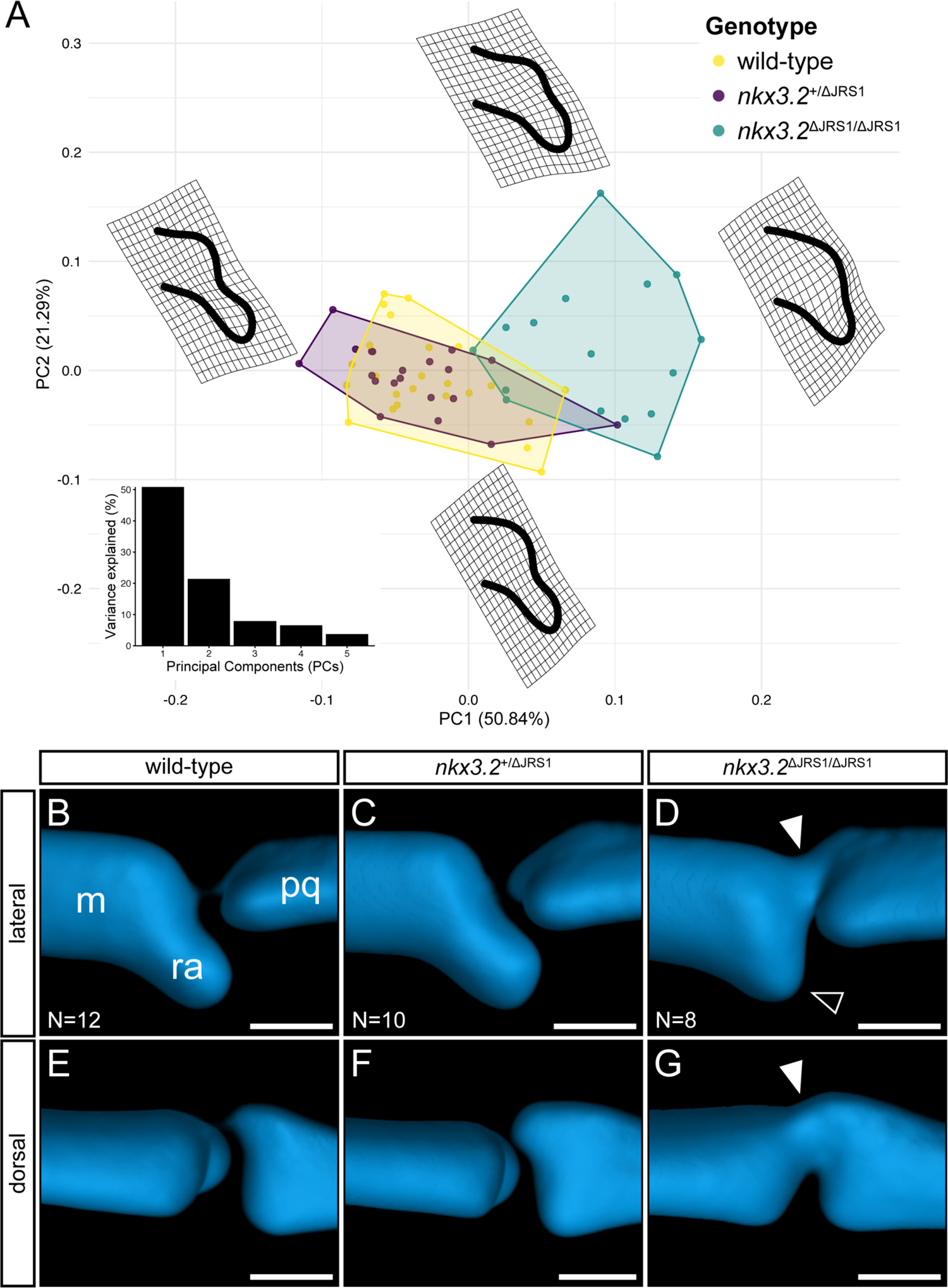
Geometric morphometric analysis of 9 dpf JRS1 deletion phenotypes. (**A**) Principal Components Analysis of geometric morphometric comparison of posterior Meckel’s cartilage shape in 9 dpf wild-type, *nkx3.2*^+/_Δ_JRS1^, and *nkx3.2*^_Δ_JRS1/_Δ_JRS1^ zebrafish. Thin-plate splines display the extremes of shape along PC1 and PC2. Inset is a histogram showing the percentage of variance explained by PC1-5. (**B-G**) Lateral and dorsal 3D renderings of the left jaw joint from averaged OPT models of 9 dpf wild-type (N=12), *nkx3.2*^+/_Δ_JRS1^ (N=10), and *nkx3.2*^_Δ_JRS1/_Δ_JRS1^ (N=8) zebrafish. White arrowheads mark the partial fusion between Meckel’s cartilage (m) and the palatoquadrate (pq). Black arrowhead marks the reduced retroarticular process (ra). Scale bars: 25 µm.

## Discussion

Previous studies have highlighted the importance of Nkx3.2 during primary jaw joint development and an overall role during chondrocyte maturation in various gnathostome species (Miller et al., 2003; Provot et al., 2006). In this study, we identified a conserved gnathostome *Nkx3.2* cis-regulatory element, *jaw joint regulatory sequence 1* (JRS1), investigated its activity in the jaw joint by generating transgenic reporter zebrafish lines, and its function by generating a CRISPR/Cas9 knockout allele (Figure 7).

**Figure 7.**
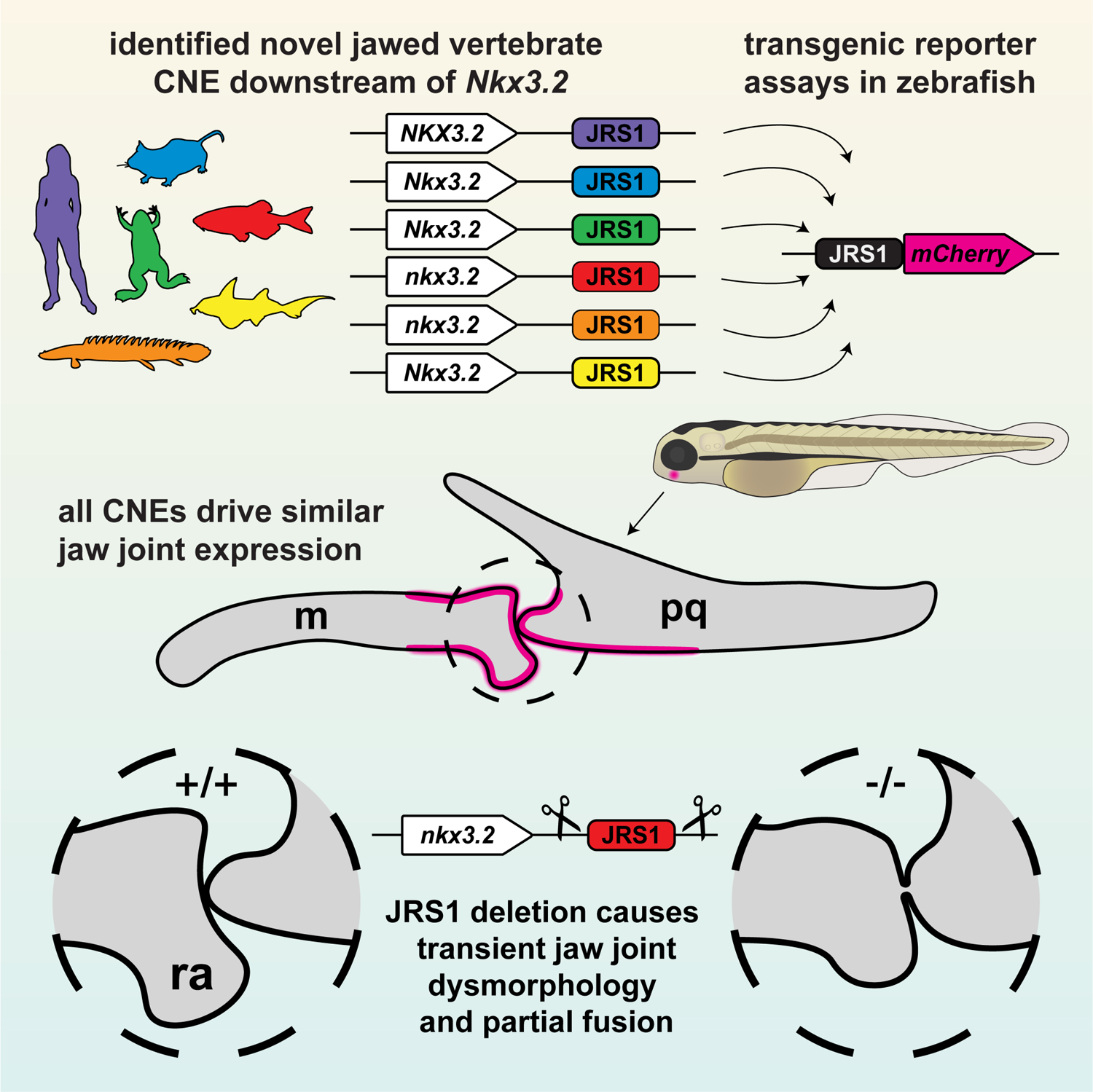
Graphical summary of this study. The conserved JRS1 enhancer was first identified as a CNE *in silico*, then confirmed with transgenesis experiments of a range of species-specific JRS1 sequences driving fluorescent reporter expression in the jaw joint. Finally, the JRS1 enhancer was knocked out in zebrafish to reveal a transient jaw joint dysmorphology and partial fusion.

We combined gene synteny analysis with non-coding sequence conservation analysis with mVISTA and detected a conserved putative *nkx3.2* enhancer sequence JRS1 in a number of gnathostome species (Figures 1, 2). Next, we applied Tol2-mediated transgenesis to test JRS1 sequences from human, mouse, frog, zebrafish, bichir, and elephant shark for enhancer activity in zebrafish. The zebrafish *nkx3.2*(JRS1):mCherry line labelled areas corresponding to the endogenous expression of *nkx3.2* in both the first pharyngeal arch and the early notochord (Video 1, Miller et al., 2003; Thisse and Thisse, 2005). We detected mCherry-labelled cells in the developing jaw joint region, including the perichondrium proximal to the joint (Figures 3, 4). *nkx3.2*(JRS1):mCherry expression overlapped with *sox10*:egfp positive neural crest derived cells at early stages, consistent with the neural crest origin of jaw joint-establishing cells (Carney et al., 2006). Notably, there was no apparent overlap of *nkx3.2*(JRS1):mCherry and *sox10*:egfp positive cells in *nkx3.2*^-/-^ mutants that displayed the fused jaw joints as early as 3 dpf, in accordance with previous report by Waldmann et al (2021). The majority of the *nkx3.2*(JRS1):mCherry-expressing cells were lining the outside of the fused jaw joint cartilages indicating that some joint or intermediate domain markers may still be present in the fusion region. This is reminiscent of the jaw joint fusion caused by the *dlx3b;4b;5a*-MO injection described by Talbot et al (2010), where *trps1*, normally expressed in the articular cartilage of the wild-type jaw joint, was detected in cells surrounding the fusion in the morphants. Despite these cell patterning changes, *nkx3.2*(JRS1):mCherry positive cells at 5 dpf displayed a similar accumulation of fluorescence reporter protein in the cell membrane of *nkx3.2*^-/-^ mutants as in wild-types, possibly indicating the absence of drastic changes in cellular processes like protein production or extracellular matrix secretion.

The zebrafish jaw joint has previously been shown to display synovial joint characteristics (Askary et al., 2016). Synovial joint development involves interzone formation after mesenchyme condensation. Interzone cells do not undergo chondrogenesis in contrast to adjacent, cartilage element-forming cells, but become flattened and densely-packed non-chondrogenic cells (Craig et al., 1987). *nkx3.2*(JRS1):mCherry expression at 3 dpf was found in cells displaying characteristic interzone cell morphology. *nkx3.2*(JRS1):mCherry expression in the mandibular arch was consistent over time, displaying a strong fluorescence signal even at 14 dpf. The scattered membranal mCherry expression we could observe from 5 dpf onwards could be a consequence of the increasing extracellular matrix production by the interzone cells, necessary to facilitate the joint cavitation process (Dowthwaite et al., 1998; Edwards et al., 1994).

Perichondrium cell-derived signalling inhibits both chondrocyte proliferation and differentiation in mouse long bone development (Alvarez et al., 2001). Our finding of JRS1 activity in the perichondrium cells lining Meckel’s cartilage and the palatoquadrate suggests the expression of *nkx3.2* in perichondrium cells during chondrocyte maturation inhibition may be important for correct shaping and sizing of the cartilaginous elements. Whether and to what extent *nkx3.2* expression in the perichondrium regulates the shape of the mandibular arch and cartilage elements more broadly requires further investigation. With a help of JRS1 deletion tests discussed further below we have started to provide some insight into this function.

Functional testing of homologous JRS1 enhancer sequences from human, mouse, frog, bichir, and elephant shark resulted in species-specific mCherry expression almost identical to what was observed in the zebrafish *nkx3.2*(JRS1):mCherry transgenic line. These data suggest that the general landscape of transcription factors binding to JRS1 is most probably conserved between gnathostomes, suggesting a conserved role for this cis-regulatory element. This is also supported by our prediction of transcription factor binding motifs broadly shared between species that are associated with functions in the pharyngeal arch patterning and skeletogenesis (Figure 2C). This list of putative transcription factor binding sites provides many candidates for validation and further research into the regulatory control of *Nkx3.2* in the pharyngeal arches.

The localisation of *Nkx3.2* to the intermediate domain of the first pharyngeal arch has been suggested as one of the key drivers of the evolution of vertebrate jaws (Cerny et al., 2010). As such, the identification of JRS1 as a jaw-joint specific enhancer of *Nkx3.2* that is conserved in gnathostomes but absent from the jawless hagfish suggests that JRS1 evolved in the gnathostome stem group after the split with Cyclostomata, and may have played a key role in the early evolution of jaws. It is also noteworthy that JRS1 appears to be functionally conserved in mammals, as the primary jaw joint region, along with its *Nkx3.2* expression domain has moved to the middle ear, forming the joint between the malleus and incus (Anthwal et al., 2013; Luo, 2007). This suggests that JRS1 may continue to be functional in driving *Nkx3.2* expression in the mammalian incudomalleolar joint.

The apparent absence of JRS1 in acanthopterygian teleost fish suggests that a single event at the base of Acanthopterygii may have resulted in the loss of JRS1, or translocation to another locus outside of the intergenic region between *nkx3.2* and *rab28*. Alternatively, JRS1 may have undergone rapid sequence evolution in acanthopterygians relative to other teleosts, and is simply too diverged to be detected based on sequence homology (Ravi and Venkatesh, 2018). It is tempting to speculate whether the apparent absence or divergence of JRS1 may be related to the evolution of the characteristic upper jaw protrusability in acanthopterygians (Lauder and Liem, 1983), as this altered functional morphology may have led to changes in selective pressures acting on the mandibular jaw joint.

The elephant shark JRS1 sequence drove reporter expression in the joint of the zebrafish first (mandibular) pharyngeal arch, consistent with other gnathostomes, but gene expression of *nkx3.2* in the related elasmobranch chondrichthyans has been reported in the intermediate domains of all pharyngeal arches, including the hyoid and gill arches (Compagnucci et al., 2013; Hirschberger et al., 2021). If we assume elephant shark *nkx3.2* gene expression mirrors that of elasmobranchs and that JRS1 drives *nkx3.2* gene expression in all pharyngeal arches, our results would support the conclusion that much of the gene regulatory landscape of osteichthyan mandibular joints is found more broadly in the pharyngeal arches of chondrichthyans (Hirschberger et al., 2021).

JRS1 knockout did not phenocopy the striking jaw joint fusion seen in zebrafish *nkx3.2* gene knockout mutants (Miyashita et al., 2020; Waldmann et al., 2021), suggesting this enhancer is not essential for *nkx3.2* expression at the time points analysed. However, as a subtle jaw joint phenotype was still evident in homozygous mutants, most notably the reduction of the RAP and partial fusion observed at 5-9 dpf, the loss of JRS1 must have some local effect on *nkx3*.2 expression, consistent with the specific jaw joint activity seen in transgenic reporter fish.

The jaw joint phenotype in homozygous JRS1 mutants appears to be rescued by approximately 14 dpf, suggesting that either *nkx3.2* expression recovers after an initial presumed decrease caused by the absence of JRS1, or that late *nkx3.2* expression is less important for shaping the developing jaw joint, and that other factors take over. At 6 dpf, we could not detect any significant differences in *nkx3.2* expression levels, supporting the former hypothesis. If this is the case, it suggests that it takes approximately one week for the phenotype to be rescued once normal *nkx3.2* expression is restored. It also indicates that JRS1 is most important to the early jaw joint expression of *nkx3.2*, prior to 6 dpf, and may contribute relatively less to later expression. Castellanos et al. (2021) recently reported subtle jaw joint phenotypes reminiscent of those seen in our homozygous JRS1 mutants in 5 dpf *hspg2* morphants that were associated with a ∼30% reduction in *nkx3.2* gene expression at 4 dpf. Their results and ours are consistent with earlier work indicating that a reduction in *nkx3.2* expression levels can result in more subtle jaw joint phenotypes, increasing in severity with increasing morpholino dosage (Miller et al., 2003).

The results of single enhancer deletions in previous studies vary widely from no phenotypic effect at all (Cunningham et al., 2018; Osterwalder et al., 2018), to subtle effects (Dickel et al., 2018), to strong effects approaching or phenocopying gene knockouts (Dobrzycki et al., 2020; Sagai et al., 2005). More severe phenotypes relative to homozygous gene knockouts likely scale with greater reductions in gene expression (Osterwalder et al., 2018), although in some cases the phenotypic effects may instead rely on an expression threshold being breached (Lam et al., 2015). It is common for developmental genes to be regulated by multiple enhancers, contributing to expression in a range of tissues and conferring a degree of functional redundancy (Chen et al., 2016; Hobert, 2010; Osterwalder et al., 2018; Wang and Goldstein, 2020). In this light, it is perhaps unsurprising that deletion of JRS1 does not completely abolish *nkx3.2* gene expression in the jaw joint, as there are likely other as-yet undiscovered enhancers that also contribute to *nkx3.2* regulation in this location in zebrafish and likely other gnathostome species as well.

This work highlights the importance of screening enhancer deletion mutants for phenotypic effects at multiple developmental stages, as the dynamic nature of gene expression by temporally-specific enhancers can mask or otherwise rescue phenotypes observed earlier in development. We encourage future studies to assess enhancer activity and gene expression to quantify these temporal dynamics, especially in known multi-enhancer systems.

In summary, we report the discovery and functional characterisation of the first described *Nkx3.2* enhancer, JRS1, with activity specific to the developing primary jaw joint. As this enhancer is conserved in sequence and activity in gnathostomes including both bony and cartilaginous fish, we conclude that it arose early in gnathostome evolution and may have been one of the earliest novel cis-regulatory elements to facilitate the evolution of jaws from the ancestral jawless state by driving the localisation of *Nkx3.2* into the jaw joint and contributing to the early jaw joint morphology. Further study of JRS1 and other as-yet undiscovered *Nkx3.2* enhancers will provide new insights into developmental regulatory network responsible for the evolution of gnathostome jaws.

## Materials and Methods

### Ethics statement

All animal experimental procedures were approved by the local ethics committee for animal research in Uppsala, Sweden (permit number C161/4 and 5.8.18-18096/2019). All procedures for the experiments were performed in accordance with the animal welfare guidelines of the Swedish National Board for Laboratory Animals.

### Conserved synteny analysis

The synteny analysis was performed on the genomic regions containing the *Nkx3.2* gene (alternative names *Nkx3-2* and *Bapx1*). Synteny data including upstream and downstream genes from *Nkx3.2* was extracted from the Ensembl database for several genomes: human, *Homo sapiens* (GRCh38.p12); mouse, *Mus musculus* (GRCm38.p6); koala, *Phascolarctos cinereus* (phaCin_unsw_v4.1); painted turtle, *Chrysemys picta* (Chrysemys_picta_bellii-3.0.3); chicken, *Gallus gallus* (GRCg6a); tropical clawed frog, *Xenopus tropicalis* (Xenopus_tropicalis_v9.1), coelacanth, *Latimeria chalumnae* (LatCha1); zebrafish, *Danio rerio* (GRCz11); spotted gar, *Lepisosteus oculatus* (LepOcu1); elephant shark, *Callorhinchus milii* (Callorhinchus_milii-6.1.3) and inshore hagfish, *Eptatretus burgeri* (Eburgeri_3.2). A genomic scaffold containing the *Nkx3.2* gene was obtained from genomic databases of the bichir, *Polypterus senegalus* (Mashima et al., 2016; Tatsumi et al., 2016).

### Conserved non-coding sequence identification and motif analysis

The genomic region surrounding the *Nkx3.2* gene, between the two immediately flanking genes (typically *Bod1l1* and *Rab28*) were collected for a number of vertebrate species and submitted to mVISTA (Frazer et al., 2004) using the default settings to search for conserved non-coding sequences. To search for JRS1 in a range of teleost species, additional sequences were extracted from the Ensembl database, representing the *nkx3.2* gene sequence and the downstream non-coding sequence until the start of the *rab28* gene. These teleost species included the arowana, *Sclerophages formosus* (fSclFor1.1); electric eel, *Electrophorus electricus* (Ee_SOAP_WITH_SSPACE); cavefish, *Astyanax mexicanus* (Astyanax_mexicanus_1.0.2); atlantic salmon, *Salmo salar* (ICSASG_v2); atlantic cod, *Gadus morhua* (gadMor3.0); soldierfish, *Myripristis murdjan* (fMyrMur1.1); seahorse, *Hippocampus comes* (H_comes_QL1_v1); tilapia, *Oreochromis niloticus* (O_niloticus_UMD_NMBU); amazon molly, *Poecilia formosa* (Poecilia_formosa-5.1.2); stickleback, *Gasterosteus aculeatus* (BROAD S1); and pufferfish, *Tetraodon nigroviridis* (TETRAODON 8.0). Non-coding sequences representing peaks of conservation were collected for human, mouse, frog, zebrafish, bichir and elephant shark and further analysed with MEME Suite (Bailey et al., 2009) to identify the conserved core of the region identified with mVISTA. Additional sequence alignment was performed using Clustal Omega (Madeira et al., 2019) and manually curated. Conserved sequence motifs in the core sequences were discovered with MEME (Bailey and Elkan, 1994) using classic discovery mode, we searched for four motifs with zero or one occurrence per sequence. Each of the discovered statistically significant motifs was matched against transcription factor binding motifs in the Vertebrate database (includes JASPAR CORE vertebrates NON-REDUNDANT (in vivo and in silico) using Tomtom (Gupta et al., 2007).

### Construct cloning

Genomic DNA *from human Homo sapiens*, mouse *Mus musculus*, frog *Xenopus tropicalis*, zebrafish *Danio rerio,* bichir *Polypterus senegalus* and elephant shark *Callorhinchus milii* was used to amplify identified JRS1 sequences. Forward primers contained four guanine residues at the 5’end, attB4 (ACAACTTTGTATAGAAAAGTT) attachment sites followed by species-specific template sequence: human 5’-GTCACACAGCTTGGAATTGGTG-3’, mouse 5’-AGTTTTACAGGTTCCTAGCCCATAC-3’, frog 5’-TCTGAACTGTTTTGCCCACATT-3’, zebrafish 5‘-AGACGTGATGCTGTGACACGCTAACTGCTG-3’, bichir 5’-GAACCGAGTGCTTTACAATTAGGTA-3’, elephant shark 5’-GAATGGAGTCACACGATAGTAATCC-3’. Reverse primers contained four guanine residues at the 5’end, attB1r (ACTGCTTTTTTGTACAAACTTG) attachment sites, adenovirus RNA polymerase II E1b minimal promoter sequence (GTGTGGAGGAGCTCAAAGTGAGGCTGAGACGCGATGGGATCCATTATATACCCTCTAGA) followed by species-specific template sequences: human 5’-AAGTGGTTCAAAGGCTAAAGTT-3’, mouse 5’-CCTCATTGCTCCACCTCTCT-3’, frog 5’-ACATTGGCACTGACAGGTAAAC-3’, zebrafish 5’-GATTTACATTTTGACGTCAAT-3’, bichir 5’-TTTCGAAATATTTGATACCGACAGT-3’, elephant shark 5’-AAAGTGCATTGTGAACAAATAGACA-3’. PCR products were subsequently recombined into pDONRP4-P1R donor vector by BP reaction according to MultiSite Gateway cloning protocol (Kwan et al., 2007; Invitrogen). For generating expression clones, entry clones created by BP reaction were used as 5’ element, PME-mCherryCAAX as middle entry clone and p3E-polyA as 3’entry clone (Kwan et al., 2007). The LR reaction was performed according to MultiSite Gateway cloning protocol (Invitrogen).

### Zebrafish transgenic lines

One-cell stage zebrafish (*Danio rerio*) embryos were microinjected with freshly mixed injection solution containing transposon expression clone (175 ng/μl) and transposase mRNA (125 ng/μl) according to Fisher et al. (2006). Injected embryos were screened at 3 dpf for mosaic expression. F0 embryos displaying the strongest mosaic *nkx3.2*(JRS1):mCherry expression were selected and raised to sexual maturity (90 dpf). Mature adults were outcrossed with AB fish. Positive *nkx3.2*(JRS1):mCherry F1 embryos were raised to establish stable transgenic lines. For generating double transgenic zebrafish, the Tg(*nkx3.2*(JRS1):mCherry) line was crossed with previously published zebrafish lines Tg(*fli1a*:EGFP) (Lawson and Weinstein, 2002) or Tg(*sox10*:egfp) (Carney et al., 2006).

For comparison between wild-type and homozygous *nkx3.2* gene mutant zebrafish, the previously reported *uu2803* null allele was crossed with double transgenic line *nkx3.2*(JRS1):mCherry/*sox10*:egfp and further incrossed to produce *nkx3.2^-/-^* fish.

### Confocal live-imaging microscopy

Confocal microscopy was performed on an inverted Leica TCS SP5 microscope using Leica Microsystem LAS-AF software. Embryos were sedated with 0.16% MS-222 and embedded in 0.8% low-melting agarose on the glass bottom of the 35mm dishes. To prevent drying, embedded embryos were covered with system water containing 0.16% MS-222. Images are presented as single images or maximum projections, as specified.

### Whole-mount in situ hybridisation

Primers were designed for zebrafish *nkx3.2* gene sequence Fw 5’-CTTCAACCACCAGCGTTATCTC-3’ and Rev 5’-ACATGTCTAGTAAACGGGCGA-3’. Fragments were cloned from zebrafish cDNA into the pCR II TOPO vector and antisense RNA probes were synthesized with either SP6 or T7 RNA polymerase and digoxigenin labelling mix (Roche). In situ hybridisation on zebrafish whole-mount 2dpf embryos was done as previously described (Filipek-Górniok et al., 2013).

### Enhancer deletion using CRISPR/Cas9

Two gRNAs targeting flanking regions of the zebrafish *nkx3.2* JRS1 enhancer were selected with the use of CRISPOR online design tool (Concordet and Haeussler, 2018): 5’-TGACGAGAGGAGCGACACGC-3’ and 5’-GCGTGTCGCTCCTCTCGTCA-3’. The gRNAs were prepared as previously described (Varshney et al., 2015). In short, annealing of the two oligos containing T7 promoter, target specific sequence, where the first two nucleotides were modified for the T7 synthesis needs, and the guide core sequence were performed. Reaction products were used as a template for the in vitro transcription (HiScribe T7 High Yield RNA Synthesis Kit, England Biolabs) and purified. Cas9 mRNA was prepared from the p-T3Ts-nCAs9 plasmid (46757 Addgene) including the restriction enzyme digestion (XbaI, New England Biolabs), in vitro transcription (mMESSAGE mMACHINE T3 Transcription Kit, Life Technologies) and product purification. Fertilized eggs were obtained by natural spawning of AB zebrafish and injected at the one-cell stage with 150 pg of *Cas9* mRNA and 50 pg of each sgRNA in RNase-free H_2_O. The efficiency of the targets was estimated by the CRISPR-STAT method (Carrington et al., 2015). Sequences of the primers used for activity testing and genotyping were 5’-GTGACACGCTAACTGCTGGA-3’ and 5’-GAACATCCTTCATGGGCTTC-3’. All primer and gRNA sequences are shown schematically in Supplementary Figure 3.

Injected F0 fish were raised to adulthood and individually outcrossed with AB zebrafish. Clutches of 5 dpf larvae were genotyped to determine the presence of the enhancer deletion in the germline of F0 parents. Three batches of 8-12 randomly selected larvae were sacrificed and lysed in a solution of 150 µL 50 µM NaOH for 20 minutes at 95°C. The lysis solution was stabilized by adding 100 µL 0.1 mM Tris. 1 µL of the resulting lysis solution was added into a 25 µL PCR reaction containing Platinum Taq DNA Polymerase and the two primers flanking the enhancer deletion site, and was run for 35 cycles. Resulting PCR products were run on 1% agarose gels to determine their size. PCR product representing enhancer deletion allele was 352 bp in length, and easily distinguishable from 790 bp wild-type allele. Four deletion-positive F0 zebrafish were outcrossed with Tg(*fli1a*:GFP) fish (Lawson and Weinstein, 2002) to produce four F1 lines, which were raised to adulthood and genotyped by fin clip. One of the four heterozygous lines (deletion allele uu3731, designated _Δ_JRS1) was selected for further analysis. These F1 *nkx3.2*^+/_Δ_JRS1^ fish were incrossed to produce F2 *nkx3.2*^_Δ_JRS1/_Δ_JRS1^ fish for analysis. JRS1 deletion was confirmed by Sanger sequencing (Eurofins Genomics) using the same PCR primers.

### Skeletal staining

Zebrafish larvae and juveniles were stained with Alcian blue and alizarin red following a protocol modified from Walker and Kimmel (2007), previously described in Waldmann et al. (2021)

### Optical Projection Tomography

A custom-built Optical Projection Tomography (OPT) (Sharpe et al., 2002; Zhang et al., 2020) system was used for imaging of 9 dpf skeletal stained zebrafish larvae. The OPT system, reconstruction algorithms, and alignment workflow were based on the previously described method (Allalou et al., 2017). All larvae were kept in 99% glycerol before they were loaded into the system for imaging. The rotational images were acquired using a 3X telecentric objective with a pixel resolution of 1.15 μm/pixel. The tomographic 3D reconstruction was done using a filtered back projection (FBP) algorithm in MATLAB (Release R2015b; MathWorks, Natick, MA) together with the ASTRA Toolbox (Palenstijn *et al*., 2013). For the data alignment, the registration toolbox elastix (Klein et al., 2010; Shamonin et al., 2014) was used. To reduce the computational time all 3D volumes in the registration were down-sampled to half the resolution.

The registration workflow was similar to the methods described by Allalou et al. (2017) where the wild-type fish were initially aligned and used to create an average reference fish using an Iterative Shape Averaging (ISA) algorithm (Rohlfing et al., 2001). All wild-type (n=12), *nkx3.2*^+/_Δ_JRS1^ (n=10), and *nkx3.2*^_Δ_JRS1/_Δ_JRS1^ (n=8) zebrafish were then aligned to the reference.

### Quantitative real-time polymerase chain reaction (qPCR)

6 dpf larvae produced from incrossing heterozygous parents were euthanised with an overdose of MS-222 (300mg/L) and the tails were removed for genotyping while the head and remaining body were stored in RNALater™ (Invitrogen). Total RNA was extracted with Trizol reagent (Fisher) from a pool of 7 larvae of each genotype, with 3 biological replicates per genotype. To prevent genomic DNA contamination, extractions were DNase-treated using the Turbo DNA-free™ kit (Ambion). cDNA was synthesised from 200ng total RNA from each sample using the SuperScript™ IV First-Strand cDNA Synthesis kit (Invitrogen) with random hexamers in a 20µL reaction. qPCR was performed on all samples in technical triplicates with PowerUp™ SYBR™ Green Master Mix using a 7500 Real Time PCR System (Applied Biosystems) and the following primers: rpl13a forward 5’-TCTGGAGGACTGTAAGAGGTATGC-3’; rpl13a reverse 5’-AGACGCACAATCTTGAGAGCAG-3’; nkx3.2 forward 5’-ACTGCGTGTCCGACTGTAACAC-3’; nkx3.2 reverse 5’-GTCTCGGTGAGTTTGAGGGA-3’. Amplicon sizes for these target genes were 148bp and 187bp, respectively. A dissociation step was performed at the end of the analysis to verify the specificity of the products, and standard curves were generated from pooled cDNA from all samples in triplicate for each target gene to verify the efficiency of the primers (R^2^>0.99). *nkx3.2* expression was normalised to *rpl13a* levels and relative quantification of gene expression was calculated using the Pfaffl method (Pfaffl, 2001), displaying the fold difference in heterozygous and homozygous mutants relative to wild-type, which was set to 1.0. FDR-adjusted P-values (Benjamini and Hochberg, 1995) are reported from pairwise Wilcoxon tests.

### Geometric Morphometric Analysis

We employed 2D geometric morphometric analysis using 100 landmarks defined along the edge of Meckel’s cartilage at the jaw joint interface. From maximum projection images generated by OPT, standardised with a lateral orientation, the shapes of Meckel’s cartilage were traced in Adobe Illustrator 2020. These shapes were then aligned by orientation and the joint interfacing surface, but not resized, before most of the anterior portion of the shapes were cut away to a standardised extent, leaving shapes representing just the posterior head of Meckel’s cartilage. These shapes were digitised using 100 equidistant landmarks using tpsDig2 (v2.3) (Rohlf, 2017), and imported into R for analysis (R Development Core Team, 2021). The two landmarks at the ends of each curve were used as fixed landmarks, and the remaining 98 as semi landmarks.

Generalized Procrustes Analysis (GPA) was used to align coordinates of the landmarks for subsequent morphospace analysis using the geomorph package (v3.3.2) (Adams et al., 2021) and bivariate plots of the PC axis were generated using the borealis package (Angelini, 2021). FDR-adjusted P-values (Benjamini and Hochberg, 1995) are reported from pairwise analyses of group means after accounting for allometric differences.

## Acknowledgments

We thank Professor Per E. Ahlberg, Dr Erika Kague, and Dr Sophie Sanchez for providing thoughtful comments on the manuscript. Professor Per E. Ahlberg also covered some lab expenses. We thank Professor Masataka Okabe at the Jikei University School of Medicine for sharing the genomic scaffold of *Polypterus senegalus*, Dr Qingming Qu for *P. senegalus* gDNA, Dr Christine Boisvert for *Callorhinchus milii* gDNA, Dr Henning Onsbring Gustafson for help with initial cloning of the zebrafish construct, Dr Mohamad Bazzi for help with geometric morphometrics, Dr Emmanouil Tsakoumis for help with RNA extraction and qPCR, and Philipp Pottmeier for access to and assistance with the 7500 Real Time PCR System. We also thank the following Erasmus students for their help during various stages of the project: Onur Özer at Bilkent University, Thibaut D’Hooge and Branco Vanhaverbeke at University College Ghent. TH was supported by Vetenskapsrådet (Starting grant 621-2012-4673). The development of the OPT system was funded by a development project at SciLifeLab Uppsala (2017) and a Technology Development grant at SciLifeLab (2018), both awarded to AA.

## Supplementary figures

**Supplementary Figure 1.**
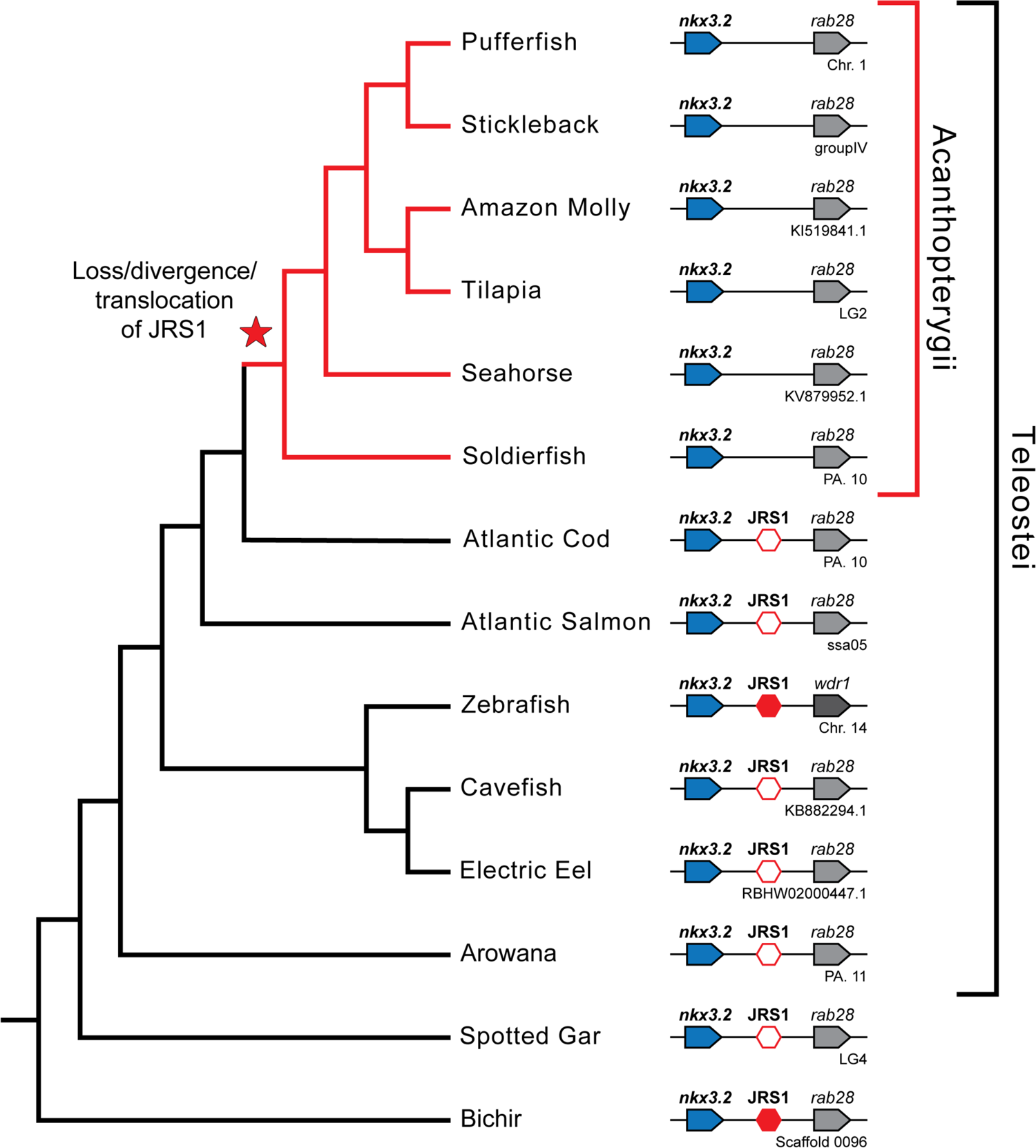
Acanthopterygian fish appear to lack JRS1. Phylogenetic tree based on Hughes et al. (2018). Pointed boxes represent gene orientation with gene names stated on top. Red hexagons mark the *Nkx3.2* enhancer (JRS1) position, where filled hexagons mark enhancers selected for *in vivo* functional characterization in this study. Below the gene order schematic of each species is marked the chromosome or contig containing this region.

**Supplementary Figure 2.**
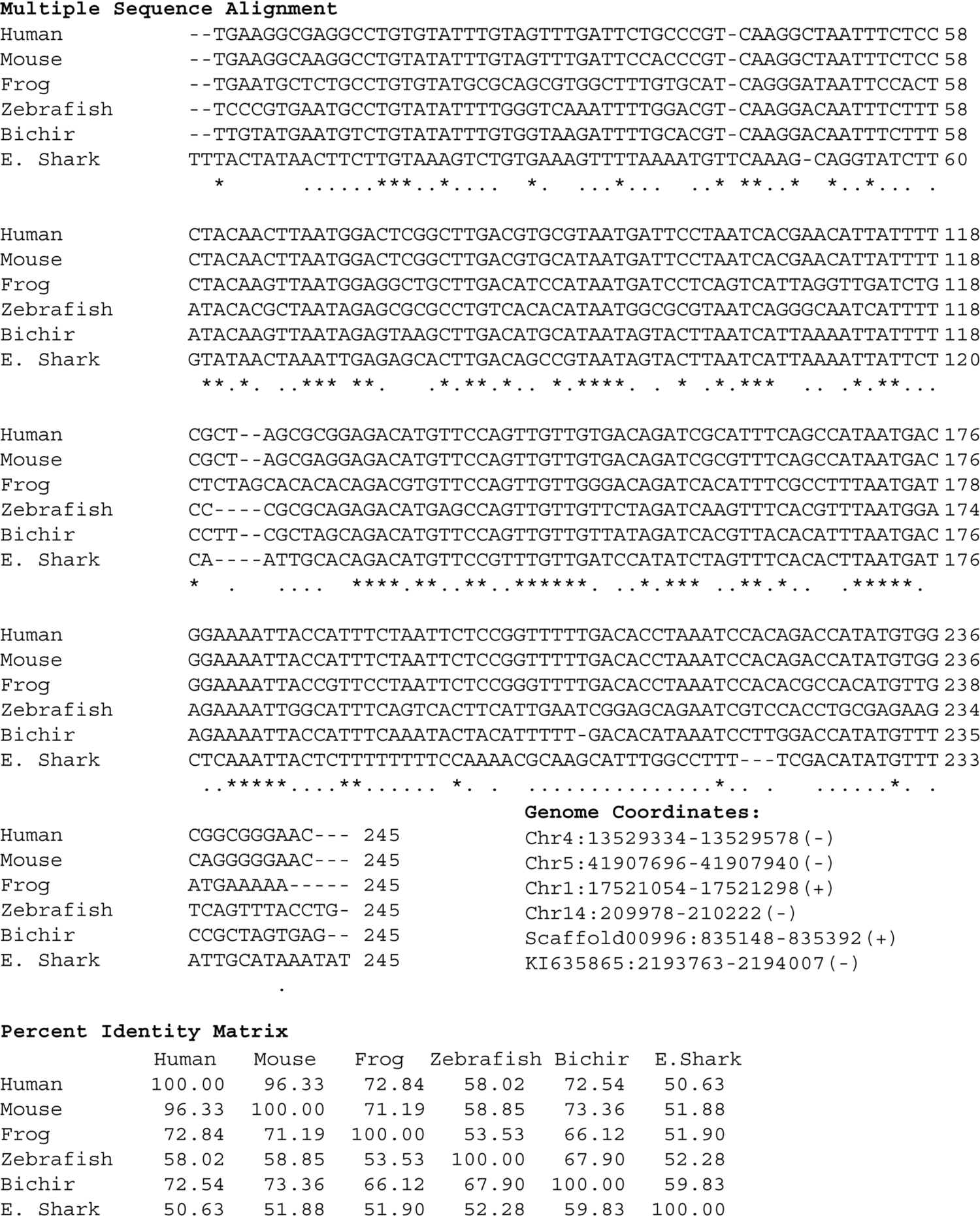
Multiple sequence alignment, genome coordinates, and percent identity matrix of the JRS1 conserved core. Asterisks mark sites in the alignment conserved in 6/6 species, dots mark sites conserved in ≥4/6 species.

**Supplementary Figure 3.**
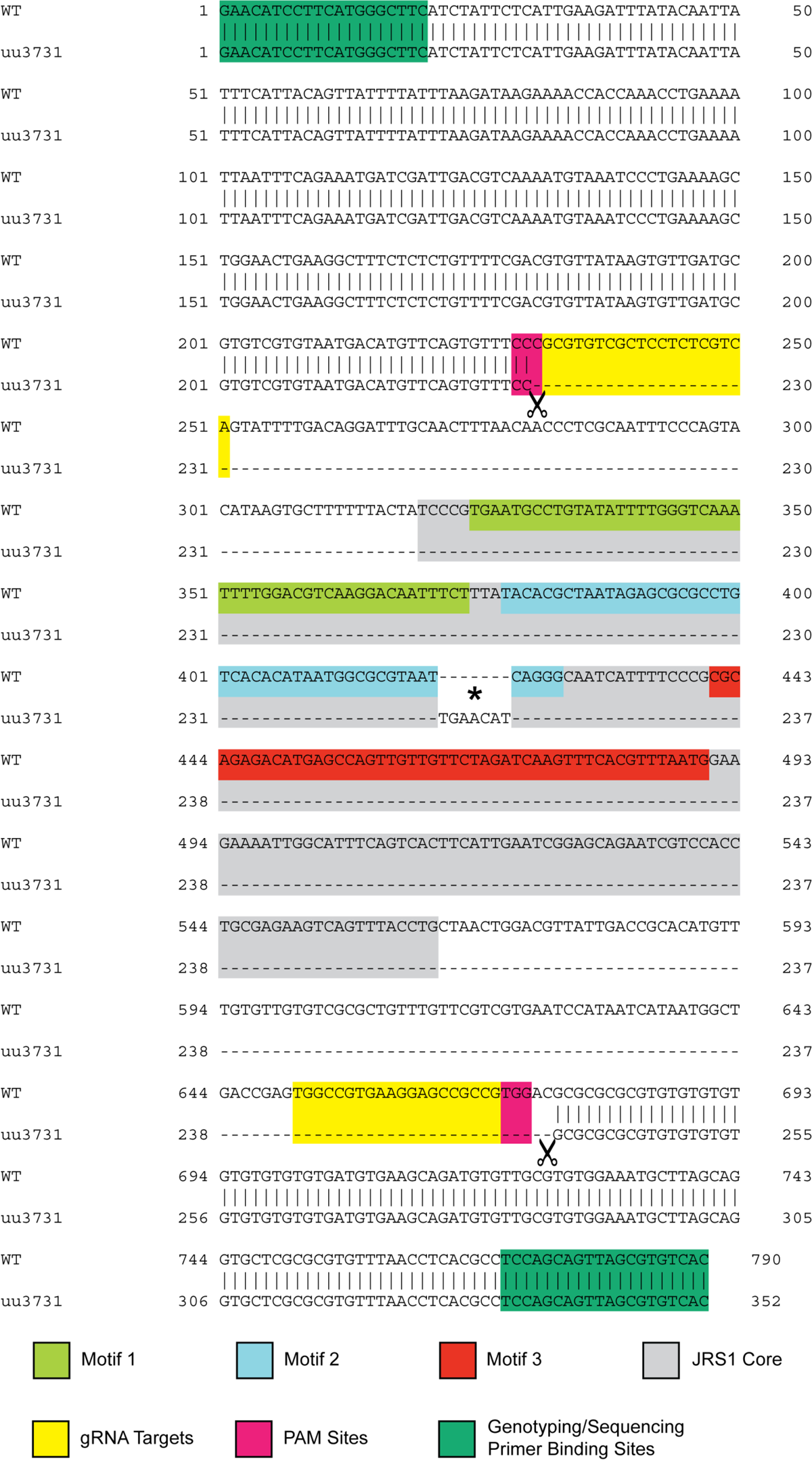
Alignment of wild-type and uu3731 JRS1 deletion alleles with annotated MEME motifs, primers, and CRISPR gRNA targets. The conserved core of the zebrafish JRS1 enhancer sequence identified by MEME is labelled in grey. Motifs 1-3 are coloured according to Figure 2B. Scissors mark the beginning and end of the deletion location, and an asterisk marks the chance insertion of a random 7bp sequence to the deletion location during DNA repair.

**Supplementary Figure 4.**
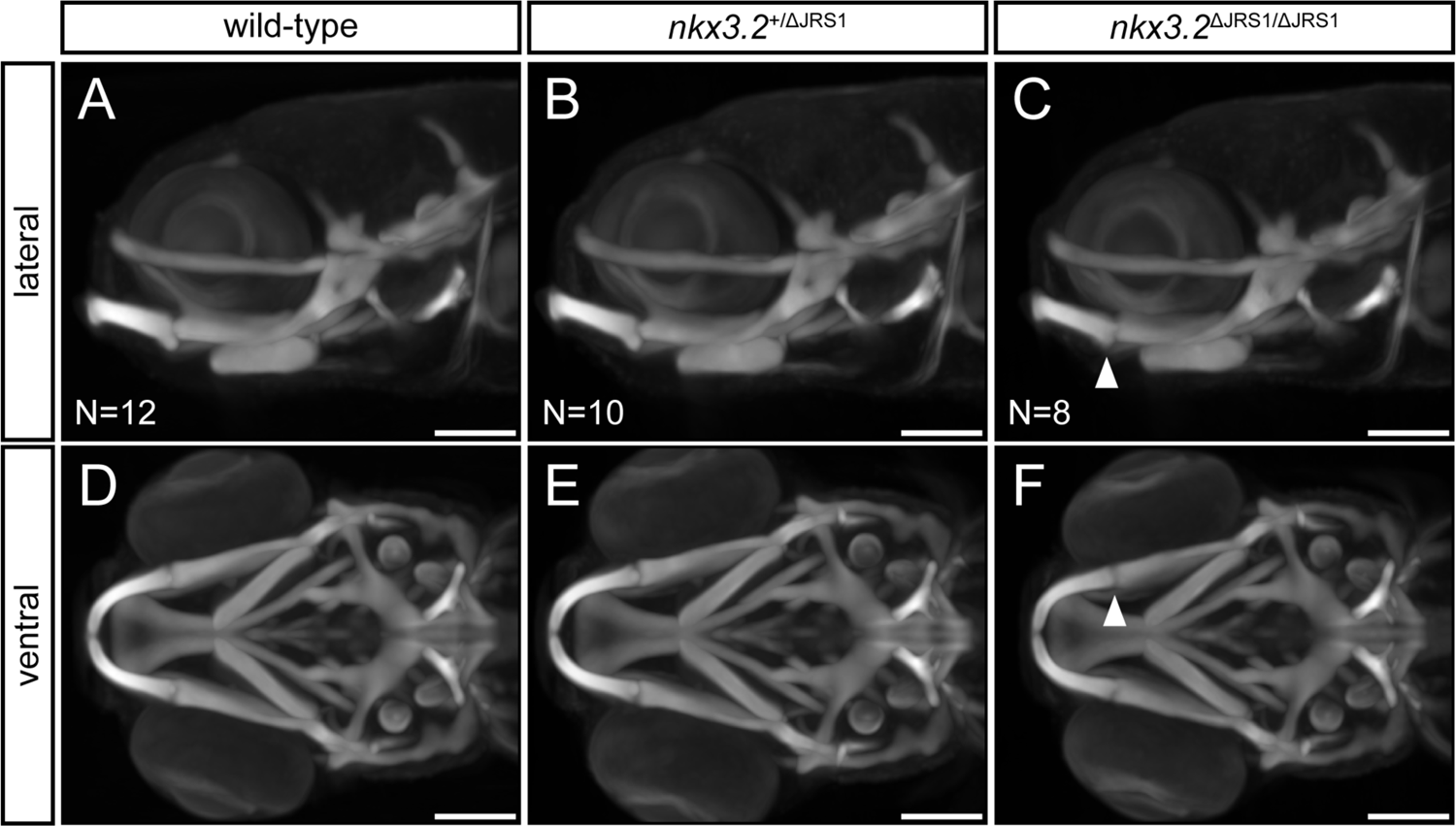
Optical Projection Tomography of 9 dpf JRS1 deletion zebrafish. (**A-C**) Lateral and (**D-F**) ventral maximum projection views of the average head skeleton of 9 dpf wild-type (N=12), *nkx3.2*^+/_Δ_JRS1^ (N=10), and *nkx3.2*^_Δ_JRS1/_Δ_JRS1^ (N=8) zebrafish. White arrowhead indicates the subtle jaw joint dysmorphology seen in *nkx3.2*^_Δ_JRS1/_Δ_JRS1^ mutants. Scale bars: 150 µM.

Figure 5K-Source Data 1 |qPCR raw values.

Figure 6A-Source Data 2 | Jaw joint landmarks TPS.

Figure 6A-Source Data 3 | Jaw joint landmark classifiers.

## References

1. Adams DC, Collyer ML, Kaliontzopoulou A, Baken EK. 2021. Geomorph: Software for geometric morphometric analyses. R package version 4.0.

2. Allalou A, Wu Y, Ghannad-Rezaie M, Eimon PM, Yanik MF. 2017. Automated deep-phenotyping of the vertebrate brain. Elife 6:e23379. doi:10.7554/eLife.23379

3. Alvarez J, Horton J, Sohn P, Serra R. 2001. The perichondrium plays an important role in mediating the effects of TGF-β1 on endochondral bone formation. Dev Dyn 221:311– 321. doi:10.1002/dvdy.1141

4. Angelini D. 2021. Tools for reproducible geometric morphometric analysis. R package version 31.03.02.

5. Anthwal N, Joshi L, Tucker AS. 2013. Evolution of the mammalian middle ear and jaw: Adaptations and novel structures. J Anat 222:147–160. doi:10.1111/j.1469-7580.2012.01526.x

6. Askary A, Smeeton J, Paul S, Schindler S, Braasch I, Ellis NA, Postlethwait J, Miller CT, Gage Crump J. 2016. Ancient origin of lubricated joints in bony vertebrates. Elife 5:e16415. doi:10.7554/eLife.16415

7. Bailey TL, Boden M, Buske FA, Frith M, Grant CE, Clementi L, Ren J, Li WW, Noble WS. 2009. MEME Suite: Tools for motif discovery and searching. Nucleic Acids Res 37:W202– W208. doi:10.1093/nar/gkp335

8. Bailey TL, Elkan C. 1994. Fitting a mixture model by expectation maximization to discover motifs in biopolymers. Proceedings Int Conf Intell Syst Mol Biol 2:28–36.

9. Benjamini Y, Hochberg Y. 1995. Controlling the False Discovery Rate: A Practical and Powerful Approach to Multiple Testing. J R Stat Soc Ser B 57:289–300. doi:10.1111/j.2517-6161.1995.tb02031.x

10. Carney TJ, Dutton KA, Greenhill E, Delfino-Machín M, Dufourcq P, Blader P, Kelsh RN. 2006. A direct role for Sox10 in specification of neural crest-derived sensory neurons. Development 133:4619–4630. doi:10.1242/dev.02668

11. Caron MMJ, Emans PJ, Surtel DAM, Van Der Kraan PM, Van Rhijn LW, Welting TJM. 2015. BAPX-1/NKX-3.2 acts as a chondrocyte hypertrophy molecular switch in osteoarthritis. Arthritis Rheumatol 67:2944–2956. doi:10.1002/art.39293

12. Carrington B, Varshney GK, Burgess SM, Sood R. 2015. CRISPR-STAT: An easy and reliable PCR-based method to evaluate target-specific sgRNA activity. Nucleic Acids Res 43:1–8. doi:10.1093/nar/gkv802

13. Castellanos BS, Reyes-Nava NG, Quintana AM. 2021. Knockdown of hspg2 is associated with mandibular jaw joint fusion and neural crest cell dysfunction in zebrafish. BMC Dev Biol 21. doi:10.21203/rs.3.rs-27670/v1

14. Cerny R, Cattell M, Sauka-Spengler T, Bronner-Fraser M, Yu F, Medeiros DM. 2010. Evidence for the prepattern/cooption model of vertebrate jaw evolution. Proc Natl Acad Sci 107:17262–17267. doi:10.1073/pnas.1009304107

15. Chang L, Yao H, Yao Z, Ho KKW, Ong MTY, Dai B, Tong W, Xu J, Qin L. 2021. Comprehensive Analysis of Key Genes, Signaling Pathways and miRNAs in Human Knee Osteoarthritis: Based on Bioinformatics. Front Pharmacol 12:1–17. doi:10.3389/fphar.2021.730587

16. Chen H, Capellini TD, Schoor M, Mortlock DP, Reddi AH, Kingsley DM. 2016. Heads, Shoulders, Elbows, Knees, and Toes: Modular Gdf5 Enhancers Control Different Joints in the Vertebrate Skeleton. PLoS Genet 12:1–27. doi:10.1371/journal.pgen.1006454

17. Cleary MA, Narcisi R, Albiero A, Jenner F, De Kroon LMG, Koevoet WJLM, Brama PAJ, Van Osch GJVM. 2017. Dynamic Regulation of TWIST1 Expression during Chondrogenic Differentiation of Human Bone Marrow-Derived Mesenchymal Stem Cells. Stem Cells Dev 26:751–761. doi:10.1089/scd.2016.0308

18. Compagnucci C, Debiais-Thibaud M, Coolen M, Fish J, Griffin JN, Bertocchini F, Minoux M, Rijli FM, Borday-Birraux V, Casane D, Mazan S, Depew MJ. 2013. Pattern and polarity in the development and evolution of the gnathostome jaw: Both conservation and heterotopy in the branchial arches of the shark, Scyliorhinus canicula. Dev Biol 377:428– 448. doi:10.1016/j.ydbio.2013.02.022

19. Concordet JP, Haeussler M. 2018. CRISPOR: Intuitive guide selection for CRISPR/Cas9 genome editing experiments and screens. Nucleic Acids Res. doi:10.1093/nar/gky354

20. Craig FM, Bentley G, Archer CW. 1987. The spatial and temporal pattern of collagens I and II and keratan sulphate in the developing chick metatarsophalangeal joint. Development 99:383–391.

21. Cunningham TJ, Lancman JJ, Berenguer M, Dong PDS, Duester G. 2018. Genomic Knockout of Two Presumed Forelimb Tbx5 Enhancers Reveals They Are Nonessential for Limb Development. Cell Rep 23:3146–3151. doi:10.1016/j.celrep.2018.05.052

22. Dickel DE, Ypsilanti AR, Pla R, Zhu Y, Barozzi I, Mannion BJ, Khin YS, Fukuda-Yuzawa Y, Plajzer-Frick I, Pickle CS, Lee EA, Harrington AN, Pham QT, Garvin TH, Kato M, Osterwalder M, Akiyama JA, Afzal V, Rubenstein JLR, Pennacchio LA, Visel A. 2018. Ultraconserved Enhancers Are Required for Normal Development. Cell 172:491–499.e15. doi:10.1016/j.cell.2017.12.017

23. Dobrzycki T, Mahony CB, Krecsmarik M, Koyunlar C, Rispoli R, Peulen-Zink J, Gussinklo K, Fedlaoui B, de Pater E, Patient R, Monteiro R. 2020. Deletion of a conserved Gata2 enhancer impairs haemogenic endothelium programming and adult Zebrafish haematopoiesis. Commun Biol 3. doi:10.1038/s42003-020-0798-3

24. Dowthwaite GP, Edwards JCW, Pitsillides AA. 1998. An essential role for the interaction between hyaluronan and hyaluronan binding proteins during joint development. J Histochem Cytochem 46:641–651. doi:10.1177/002215549804600509

25. Edwards JC, Wilkinson LS, Jones HM, Soothill P, Henderson KJ, Worrall JG, Pitsillides AA. 1994. The formation of human synovial joint cavities: a possible role for hyaluronan and CD44 in altered interzone cohesion. J Anat 185:355–67.

26. Filipek-Górniok B, Holmborn K, Haitina T, Habicher J, Oliveira MB, Hellgren C, Eriksson I, Kjellén L, Kreuger J, Ledin J. 2013. Expression of chondroitin/dermatan sulfate glycosyltransferases during early zebrafish development. Dev Dyn 242:964–975. doi:10.1002/dvdy.23981

27. Fisher S, Grice EA, Vinton RM, Bessling SL, Urasaki A, Kawakami K, McCallion AS. 2006. Evaluating the biological relevance of putative enhancers using Tol2 transposon-mediated transgenesis in zebrafish. Nat Protoc 1:1297–1305. doi:10.1038/nprot.2006.230

28. Frazer KA, Pachter L, Poliakov A, Rubin EM, Dubchak I. 2004. VISTA: Computational tools for comparative genomics. Nucleic Acids Res 32:W273–W279. doi:10.1093/nar/gkh458

29. Germanguz I, Gitelman I. 2012. All four twist genes of zebrafish have partially redundant, but essential, roles in patterning the craniofacial skeleton. J Appl Ichthyol 28:364–371. doi:10.1111/j.1439-0426.2012.02016.x

30. Guo C, Sun Y, Zhou B, Adam RM, Li XK, Pu WT, Morrow BE, Moon A, Li X. 2011. A Tbx1-Six1/Eya1-Fgf8 genetic pathway controls mammalian cardiovascular and craniofacial morphogenesis. J Clin Invest 121:1585–1595. doi:10.1172/JCI44630

31. Gupta S, Stamatoyannopoulos JA, Bailey TL, Noble W. 2007. Quantifying similarity between motifs. Genome Biol 8:R24. doi:10.1186/gb-2007-8-2-r24

32. Hasei J, Teramura T, Takehara T, Onodera Y, Horii T, Olmer M, Hatada I, Fukuda K, Ozaki T, Lotz MK, Asahara H. 2017. TWIST1 induces MMP3 expression through up-regulating DNA hydroxymethylation and promotes catabolic responses in human chondrocytes. Sci Rep 7:1–10. doi:10.1038/srep42990

33. Hirschberger C, Sleight VA, Criswell KE, Clark SJ, Gillis A. 2021. Conserved and unique transcriptional features of pharyngeal arches in the skate (Leucoraja erinacea) and evolution of the jaw. Mol Biol Evol. doi: https://doi.org/10.1093/molbev/msab123

34. Hissnauer TN, Baranowsky A, Pestka JM, Streichert T, Wiegandt K, Goepfert C, Beil FT, Albers J, Schulze J, Ueblacker P, Petersen JP, Schinke T, Meenen NM, Pörtner R, Amling M. 2010. Identification of molecular markers for articular cartilage. Osteoarthr Cartil 18:1630–1638. doi:10.1016/j.joca.2010.10.002

35. Hobert O. 2010. Gene regulation: Enhancers stepping out of the shadow. Curr Biol 20:R697– R699. doi:10.1016/j.cub.2010.07.035

36. Hughes LC, Ortí G, Huang Y, Sun Y, Baldwin CC, Thompson AW, Arcila D, Betancur R, Li C, Becker L, Bellora N, Zhao X, Li X, Wang M, Fang C, Xie B, Zhoui Z, Huang H, Chen S, Venkatesh B, Shi Q. 2018. Comprehensive phylogeny of ray-finned fishes (Actinopterygii) based on transcriptomic and genomic data. Proc Natl Acad Sci U S A 115:6249–6254. doi:10.1073/pnas.1719358115

37. Jezewski PA, Fang PK, Payne-Ferreira TL, Yelick PC. 2009. Alternative splicing, phylogenetic analysis, and craniofacial expression of zebrafish tbx22. Dev Dyn 238:1605–1612. doi:10.1002/dvdy.21962

38. Kim Y Il, No Lee J, Bhandari S, Nam IK, Yoo KW, Kim SJ, Oh GS, Kim HJ, So HS, Choe SK, Park R. 2015. Cartilage development requires the function of Estrogen-related receptor alpha that directly regulates sox9 expression in zebrafish. Sci Rep 5:1–10. doi:10.1038/srep18011

39. Klein S, Staring M, Murphy K, Viergever MA, Pluim JPW. 2010. Elastix: A toolbox for intensity-based medical image registration. IEEE Trans Med Imaging 29:196–205. doi:10.1109/TMI.2009.2035616

40. Kwan KM, Fujimoto E, Grabher C, Mangum BD, Hardy ME, Campbell DS, Parant JM, Yost HJ, Kanki JP, Chien C Bin. 2007. The Tol2kit: A multisite gateway-based construction Kit for Tol2 transposon transgenesis constructs. Dev Dyn 236:3088–3099. doi:10.1002/dvdy.21343

41. Lam DD, de Souza FSJ, Nasif S, Yamashita M, López-Leal R, Otero-Corchon V, Meece K, Sampath H, Mercer AJ, Wardlaw SL, Rubinstein M, Low MJ. 2015. Partially Redundant Enhancers Cooperatively Maintain Mammalian Pomc Expression Above a Critical Functional Threshold. PLoS Genet 11:1–21. doi:10.1371/journal.pgen.1004935

42. Lauder G V, Liem KF. 1983. Comparative Zoology. Bull Museum Comp Zool 150:95–197. doi:10.1038/221101b0

43. Lawson ND, Weinstein BM. 2002. In vivo imaging of embryonic vascular development using transgenic zebrafish. Dev Biol 248:307–318. doi:10.1006/dbio.2002.0711

44. Lukas P, Olsson L. 2018a. Bapx1 is required for jaw joint development in amphibians. Evol Dev 20:192–206. doi:10.1111/ede.12267

45. Lukas P, Olsson L. 2018b. Bapx1 upregulation is associated with ectopic mandibular cartilage development in amphibians. Zool Lett 4:16. doi:10.1186/s40851-018-0101-3

46. Luo ZX. 2007. Transformation and diversification in early mammal evolution. Nature 450:1011–1019. doi:10.1038/nature06277

47. Machon O, Masek J, Machonova O, Krauss S, Kozmik Z. 2015. Meis2 is essential for cranial and cardiac neural crest development. BMC Dev Biol 15:1–16. doi:10.1186/s12861-015-0093-6

48. Madeira F, Park YM, Lee J, Buso N, Gur T, Madhusoodanan N, Basutkar P, Tivey ARN, Potter SC, Finn RD, Lopez R. 2019. The EMBL-EBI search and sequence analysis tools APIs in 2019. Nucleic Acids Res 47:W636–W641. doi:10.1093/nar/gkz268

49. Mashima J, Kodama Y, Kosuge T, Fujisawa T, Katayama T, Nagasaki H, Okuda Y, Kaminuma E, Ogasawara O, Okubo K, Nakamura Y, Takagi T. 2016. DNA data bank of Japan (DDBJ) progress report. Nucleic Acids Res 44:D51–D57. doi:10.1093/nar/gkv1105

50. Mcadams HH, Arkin A. 1997. Stochastic mechanisms in gene expression. Proc Natl Acad Sci U S A 94:814–819. doi:10.1073/pnas.94.3.814

51. Melvin VS, Feng W, Hernandez-Lagunas L, Artinger KB, Williams T. 2013. A morpholino-based screen to identify novel genes involved in craniofacial morphogenesis. Dev Dyn 242:817–831. doi:10.1002/dvdy.23969

52. Miller CT, Yelon D, Stainier DYR, Kimmel CB. 2003. Two endothelin 1 effectors, hand2 and bapx1, pattern ventral pharyngeal cartilage and the jaw joint. Development 130:1353–1365. doi:10.1242/dev.00339

53. Miyashita T. 2018. Development, Anatomy, and Phylogenetic Relationships of Jawless Vertebrates and Tests of Hypotheses about Early Vertebrate Evolution. University of Alberta.

54. Miyashita T, Baddam P, Smeeton J, Oel AP, Natarajan N, Gordon B, Palmer AR, Crump JG, Graf D, Allison WT. 2020. nkx3.2 mutant zebrafish accommodate the jaw joint loss through a phenocopy of the head shapes of Paleozoic jawless fish. J Exp Biol 223:jeb216945. doi:10.1242/jeb.216945

55. Oh HK, Park M, Choi SW, Jeong DU, Kim BJ, Kim JA, Choi HJ, Lee J, Cho Y, Kim JH, Seong JK, Choi BH, Min BH, Kim DW. 2021. Suppression of Osteoarthritis progression by post-natal Induction of Nkx3.2. Biochem Biophys Res Commun 571:188–194. doi:10.1016/j.bbrc.2021.07.074

56. Osterwalder M, Barozzi I, Tissiéres V, Fukuda-Yuzawa Y, Mannion BJ, Afzal SY, Lee EA, Zhu Y, Plajzer-Frick I, Pickle CS, Kato M, Garvin TH, Pham QT, Harrington AN, Akiyama JA, Afzal V, Lopez-Rios J, Dickel DE, Visel A, Pennacchio LA. 2018. Enhancer redundancy provides phenotypic robustness in mammalian development. Nature 554:239–243. doi:10.1038/nature25461

57. Palenstijn WJ, Batenburg KJ, Sijbers J. 2013. The ASTRA Tomography ToolboxProceedings of the 13th International Conference on Computational and Mathematical Methods in Science and Engineering. pp. 1139–1145.

58. Papaioannou VE. 2014. The t-box gene family: Emerging roles in development, Stem cells and cancer. Development 141:3819–3833. doi:10.1242/dev.104471

59. Pfaffl MW. 2001. A new mathematical model for relative quantification in real-time RT–PCR. Nucleic Acids Res 29:E45. doi: https://doi.org/10.1093/nar/29.9.e45

60. Piotrowski T, Ahn DG, Schilling TF, Nair S, Ruvinsky I, Geisler R, Rauch GJ, Haffter P, Zon LI, Zhou Y, Foott H, Dawid IB, Ho RK. 2003. The zebrafish van gogh mutation disrupts tbx1, which is involved in the DiGeorge deletion syndrome in humans. Development 130:5043–5052. doi:10.1242/dev.00704

61. Provot S, Kempf H, Murtaugh LC, Chung U, Kim DW, Chyung J, Kronenberg HM, Lassar AB. 2006. Nkx3.2/Bapx1 acts as a negative regulator of chondrocyte maturation. Development 133:651–62. doi:10.1242/dev.02258

62. R Development Core Team R. 2021. R: A Language and Environment for Statistical Computing.

63. Ravi V, Venkatesh B. 2018. The Divergent Genomes of Teleosts. Annu Rev Anim Biosci 6:47– 68. doi:10.1146/annurev-animal-030117-014821

64. Reinhold MI, Kapadia RM, Liao Z, Naski MC. 2006. The Wnt-inducible transcription factor Twist1 inhibits chondrogenesis. J Biol Chem 281:1381–1388. doi:10.1074/jbc.M504875200

65. Rohlf FJ. 2017. Tpsdig2, digitise landmarks and outlines, version 2.3. Stony Brook, NY: Department of Ecology and Evolution, State University of New York at Stony Brook.

66. Rohlfing T, Brandt R, Maurer CR, Menzel R. 2001. Bee brains, B-splines and computational democracy: Generating an average shape atlasProceedings of the Workshop on Mathematical Methods in Biomedical Image Analysis. IEEE. pp. 187–194. doi:10.1109/mmbia.2001.991733

67. Sagai T, Hosoya M, Mizushina Y, Tamura M, Shiroishi T. 2005. Elimination of a long-range cis-regulatory module causes complete loss of limb-specific Shh expression and truncation of the mouse limb. Development 132:797–803. doi:10.1242/dev.01613

68. Schilling TF, Kimmel CB. 1994. Segment and cell type lineage restrictions during pharyngeal arch development in the zebrafish embryo. Development 120:483–494.

69. Shamonin DP, Bron EE, Lelieveldt BPF, Smits M, Klein S, Staring M. 2014. Fast parallel image registration on CPU and GPU for diagnostic classification of Alzheimer’s disease. Front Neuroinform 7:50. doi:10.3389/fninf.2013.00050

70. Sharpe J, Ahlgren U, Perry P, Hill B, Ross A, Hecksher-Sørensen J, Baldock R, Davidson D. 2002. Optical projection tomography as a tool for 3D microscopy and gene expression studies. Science 296:541–545. doi:10.1126/science.1068206

71. Son YO, Park S, Kwak JS, Won Y, Choi WS, Rhee J, Chun CH, Ryu JH, Kim DK, Choi HS, Chun JS. 2017. Estrogen-related receptor γ causes osteoarthritis by upregulating extracellular matrix-degrading enzymes. Nat Commun 8. doi:10.1038/s41467-017-01868-8

72. Square T, Jandzik D, Cattell M, Coe A, Doherty J, Medeiros DM. 2015. A gene expression map of the larval Xenopus laevis head reveals developmental changes underlying the evolution of new skeletal elements. Dev Biol 397:293–304. doi:10.1016/j.ydbio.2014.10.016

73. Swartz ME, Sheehan-Rooney K, Dixon MJ, Eberhart JK. 2011. Examination of a palatogenic gene program in zebrafish. Dev Dyn 240:2204–2220. doi:10.1002/dvdy.22713

74. Takai H, van Wijnen AJ, Ogata Y. 2019. Induction of chondrogenic or mesenchymal stem cells from human periodontal ligament cells through inhibition of Twist2 or Klf12. J Oral Sci 61:313–320. doi:10.2334/josnusd.18-0224

75. Tang J, Liu T, Wen X, Zhou Z, Yan J, Gao J, Zuo J. 2021. Estrogen-related receptors: novel potential regulators of osteoarthritis pathogenesis. Mol Med 27. doi:10.1186/s10020-021-00270-x

76. Tatsumi N, Kobayashi R, Yano T, Noda M, Fujimura K, Okada N, Okabe M. 2016. Molecular developmental mechanism in polypterid fish provides insight into the origin of vertebrate lungs. Sci Rep 6:1–10. doi:10.1038/srep30580

77. Tavares ALP, Cox TC, Maxson RM, Ford HL, Clouthier DE. 2017. Negative regulation of endothelin signaling by SIX1 is required for proper maxillary development. Dev 144:2021–2031. doi:10.1242/dev.145144

78. Thisse C, Thisse B. 2005. High Throughput Expression Anlysis of ZF-Models Consortium clones. ZFIN Direct Data Submiss (http://zfin.org).

79. Tucker AS, Watson RP, Lettice LA, Yamada G, Hill RE. 2004. Bapx1 regulates patterning in the middle ear: altered regulatory role in the transition from the proximal jaw during vertebrate evolution. Development 131:1235–1245. doi:10.1242/dev.01017

80. Varshney GK, Pei W, Lafave MC, Idol J, Xu L, Gallardo V, Carrington B, Bishop K, Jones M, Li M, Harper U, Huang SC, Prakash A, Chen W, Sood R, Ledin J, Burgess SM. 2015. High-throughput gene targeting and phenotyping in zebrafish using CRISPR/Cas9. Genome Res 25:1030–1042. doi:10.1101/gr.186379.114

81. Waldmann L, Leyhr J, Zhang H, Öhman-Mägi C, Allalou A, Haitina T. 2021. The Broad Role of Nkx3.2 in the Development of the Zebrafish Axial Skeleton. PLoS One 16:e0255953. doi:10.1371/journal.pone.0255953

82. Walker MB, Kimmel CB. 2007. A two-color acid-free cartilage and bone stain for zebrafish larvae. Biotech Histochem 82:23–28. doi:10.1080/10520290701333558

83. Wang X, Goldstein DB. 2020. Enhancer Domains Predict Gene Pathogenicity and Inform Gene Discovery in Complex Disease. Am J Hum Genet 106:215–233. doi:10.1016/j.ajhg.2020.01.012

84. Wigner NA, Soung DY, Einhorn TA, Drissi H, Gerstenfeld LC. 2013. Functional role of Runx3 in the regulation of aggrecan expression during cartilage development. J Cell Physiol 228:2232–2242. doi:10.1002/jcp.24396

85. Yano F, Ohba S, Murahashi Y, Tanaka S, Saito T, Chung U il. 2019. Runx1 contributes to articular cartilage maintenance by enhancement of cartilage matrix production and suppression of hypertrophic differentiation. Sci Rep 9:1–9. doi:10.1038/s41598-019-43948-3

86. Zhang H, Waldmann L, Manuel R, Boije H, Haitina T, Allalou A. 2020. zOPT: an open source optical projection tomography system and methods for rapid 3D zebrafish imaging. Biomed Opt Express 11:4290. doi:10.1364/boe.393519

87. Zhu M, Yu X, Ahlberg PE, Choo B, Lu J, Qiao T, Qu Q, Zhao W, Jia L, Blom H, Zhu Y. 2013. A Silurian placoderm with osteichthyan-like marginal jaw bones. Nature 502:188–193. doi:10.1038/nature12617

88. Zuniga E, Stellabotte F, Gage Crump J. 2010. Jagged-Notch signaling ensures dorsal skeletal identity in the vertebrate face. Development 137:1843–1852. doi:10.1242/dev.049056

